# Maternal lipids prime quiescent neural stem cells to reactivate in response to dietary nutrients

**DOI:** 10.64898/2026.01.05.697812

**Authors:** Md Ausrafuggaman Nahid, Susan E. Doyle, Kelly E. Dunham, Michelle L. Bland, Sarah E. Siegrist

## Abstract

Stem cell quiescence--a state of mitotic and metabolic dormancy--is essential for tissue homeostasis and for coordinating growth with nutrient availability. Switching between quiescence and proliferation is controlled through stem cell intrinsic and extrinsic cues, with diet being key as diet provides macro-and micro-nutrients needed for synthesizing new membrane, protein, and nucleic acids. Yet, it remains unclear what nutrients control the stem cell switch and whether nutrient sources other than diet are required. Here, we report that lipids deposited maternally in the embryo regulate reactivation of *Drosophila* neural stem cells (neuroblasts) from quiescence. This maternal nutrient source is in addition to the known dietary amino acids required during larval feeding^1^. Females fed reduced lipid diets or carrying mutations in genes essential for lipid deposition and metabolism produce larvae with fewer stored lipids in neural tissues. Reduced neural lipid stores result in delayed glial growth and neuroblast reactivation due to the inability of neuroblasts to activate PI3-kinase signaling in response to diet-induced expression of insulin-like peptides. Thus, neuroblasts rely on two nutrient sources, maternal and dietary, raising the possibility that quiescent stem cells in general access and utilize stored and acquired nutrients coordinately to switch between stem cell states.

---

Maternal deposition of RNA, protein, lipid, and glycogen is essential for supporting development of the fertilized egg^2–5^. Once eggs implant into the uterus in mammals or hatch in other species, continued growth and development relies on the acquisition of outside nutrients. A relatively simple, yet functional CNS is present at hatching in many animal species including fish, amphibians and insect larvae, that contains both neural circuits that control feeding behavior and a diverse array of neural stem cell and progenitor cell types. The transition from non-feeding to feeding stages marks a critical period in an organism’s life cycle as egg-supplied nutrients become depleted. Importantly, during this transition, most cell divisions are halted organism-wide and resume only after outside nutrients are acquired^1,6,7^. At the organism level, dietary nutrients are processed in the gut and delivered to cells in peripheral tissues providing the macromolecules needed to support synthesis of new nucleic acids, proteins, lipids, and essential metabolites^8,9^. In the CNS, once nutrients are sensed and received, neural stem cells and progenitors reinitiate cell divisions, generating new progeny to support the ongoing expansion and diversification of the CNS^1,10,11^. In *Drosophila* larvae, the current model posits that peripheral tissues send a triggering mitogen to the brain in response to animal feeding^1,12,13^. This leads to the synthesis and release of Dilp-2 from a group of neurosecretory neurons in the brain and possibly other Dilps from other sources including glia^12–16^. Dilp-2, one of eight *Drosophila* insulin-like peptides, is regulated by nutrient availability and during favorable conditions (presence of amino acids) is released systemically into the hemolymph and locally into the brain^17,18^. Dilp-2 binds the Insulin-like tyrosine kinase receptor in both neuroblasts and surrounding glia, leading to downstream activation of PI3-kinase (Phosphoinositide 3-kinase), Akt and TOR. This leads to cellular uptake of glucose, synthesis of new protein and lipid, and changes in cell cycle gene expression. Remarkedly, many stem cells found in other tissue types in other species across the animal kingdom control the switch from quiescence to proliferation using the same growth signaling pathways, including intestinal and neural stem cells and even the blastocyst itself^6,19–22^.

## Quiescent neuroblasts utilize brain LDs to reactivate

Using *Drosophila* neuroblasts as a model, we set out to better understand how nutrients control the stem cell switch from quiescence to proliferation. We investigated the role of lipids by assaying lipid droplets (LDs). LDs are organelles that store neutral lipids as triacylglycerol (TAG) and sterol esters, and during feeding stages, accumulate in fat body adipocytes and in glia in neural tissue^23–27^. During non-feeding stages or under nutrient deprivation, LDs are consumed to meet energy demands for survival and continued development. LDs also accumulate in dauer-stage *C. elegans* and in quiescent mammalian neural stem cells where they play a role in controlling stem cell behavior^28–31^. At freshly hatched (FH) larval stages (0 hours ALH), before feeding, brain LD number was more than two-fold higher compared to 24 hours later, after animals fed on a complete Bloomington (BL) diet (Fig. 1A–C and SF1S). Brain LD number was further reduced in animals maintained on PBS for 24 hours, indicating that LD mobilization is accelerated under nutrient deprivation (Fig. 1C). At the FH stage, LDs were distributed throughout the brain, in glia, neurons, quiescent neuroblasts, and in the proliferating mushroom body (MB) neuroblasts (SF1A-C). At 24 hours after feeding, LDs were mostly absent in these cell types (SF1D-F). Next, we examined brain LDs in two types of mutants: those with defects in LD allocation during embryogenesis and those with defects in LD breakdown. *Jabba* and *klar* (klarsicht) mutants misallocate LDs to the yolk, while *Lsd-2* (Lipid storage droplet-2) mutants maintain a smaller pool; in all cases, fewer LDs reach the embryo proper^2,32–36^. At FH larval stages and 24 hours after feeding, *Jabba*, *Lsd-2*, and *klar* mutants had fewer LDs in their brains compared to wild type Oregon R (OR) animals (Fig. 1D,E,H,I and SF1J). In contrast, mutants for the TAG lipase *bmm* (*brummer*, mammalian ATGL^37^) had more brain LDs (Fig. 1F-I, SF1K). Next, we assayed neuroblast reactivation from quiescence. In wild-type OR animals at FH stages, only the four MB and one VL (ventrolateral) neuroblast are dividing (Fig. 1J,L and SF2A,B)^1,38,39^. After 24 hours of BL feeding, approximately sixty percent of quiescent neuroblasts identified based on expression of the Deadpan (Dpn) transcription factor reactivated and re-entered cell cycle, based on their incorporation of the thymidine analogue EdU (Fig. 1K,L,Q and SF2C). In contrast, all animals with mutations in genes controlling LD allocation or breakdown showed fewer reactivated neuroblasts (Fig. 1M-Q, SF1L, and SF2D). After 48 hours of BL feeding, most neuroblasts in mutant animals reactivated (SF1M-R). We conclude that LDs are consumed and utilized to reactivate quiescent neuroblasts.

**Figure 1:**
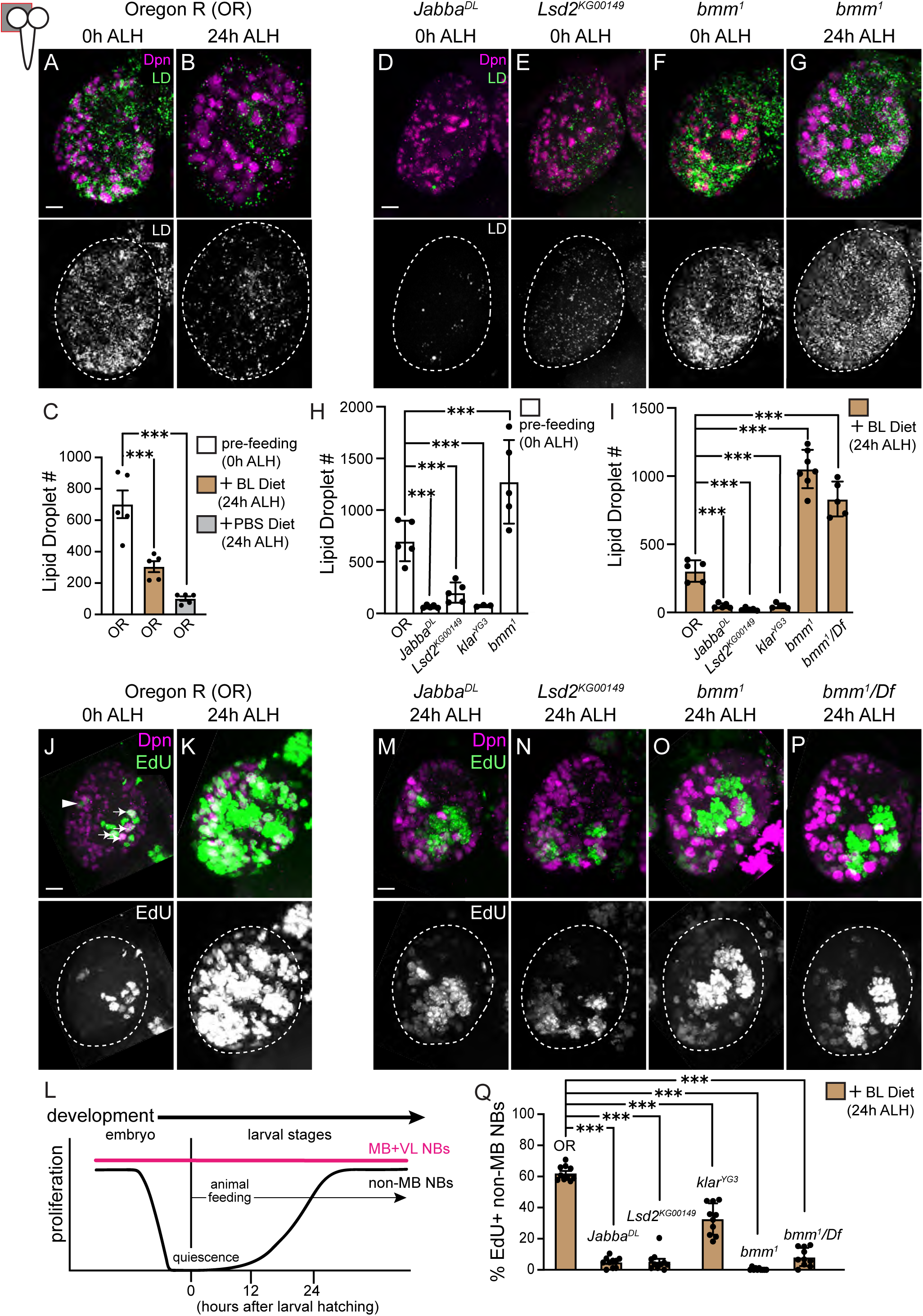
Lipid droplet consumption correlates with neuroblast reactivation. (A,B,D-G) Maximum intensity projections (MIPs) of single brain hemispheres. Cartoon upper left depicts image location (gray highlight) in panels: anterior is to the top, and midline, right in this and all subsequent figures. Time points and genotypes listed above and markers within panels. Dpn (Deadpan), a transcription factor, labels neuroblasts and Nile Red, Lipid droplets (LD). Top panels, colored overlays with grayscale images below with brain hemispheres outlined. (C,H,I) Quantification of LD number in Oregon R and mutant animals at different times and different conditions: a white colored column in this and all subsequent histograms denotes no food, tan denotes Bloomington food (BL), and gray, Phosphate buffered saline (PBS) only. (J,K,M-P) MIPs of single brain hemispheres after EdU feeding for 3 hours (J) or 24 hours on BL food (K,M-P) with quantification (Q). Top panels, colored overlays with grayscale images below. (L) Timeline of neuroblast entry into quiescence and exit from quiescence in response to BL feeding. MB (mushroom body, four per brain hemisphere, arrows in J) and the single VL (ventro-lateral, arrowhead in J) neuroblasts divide continually during the embryonic to larval transition. Each data point (C,H,I,Q) represents one brain hemisphere from one animal. Mean ± SD; one-way ANOVA with Dunnett’s post hoc test versus control (***p≤0.05). Scale bar in (A,D,J,M) equals 10μm; panels to the right in the same row share the same scale in this and all subsequent figures. Panel by panel genotypes listed in Supplemental Table 1.

Next, we tested whether LDs in the brain at FH stages support neuroblast reactivation, or whether peripheral LDs are required. Bmm associates with LDs in both the brain and the peripheral fat body at FH stages and 24 hours after feeding (SF1G-I). We used GAL4/UAS to knock down *bmm* and block lipolysis in a cell- and tissue-type specific manner. At 24 hours after BL feeding, animals expressing *UAS-bmmRNAi* in neuroblasts (*worGAL4*), glia (*repoGAL4*), or cortex glia (*NP0577GAL4*), a glial subset that ensheath neuroblasts and their newborn progeny, had reduced neuroblast reactivation compared to controls expressing GAL4 alone (Fig. 2A,B,D,F-H,J). In contrast, animals expressing *UAS-bmmRNAi* in the peripheral lipid storage tissue, fat body (*r4GAL4*) showed little to no difference compared to control (Fig. 2E). We then assayed LD number in knockdown animals.

**Figure 2:**
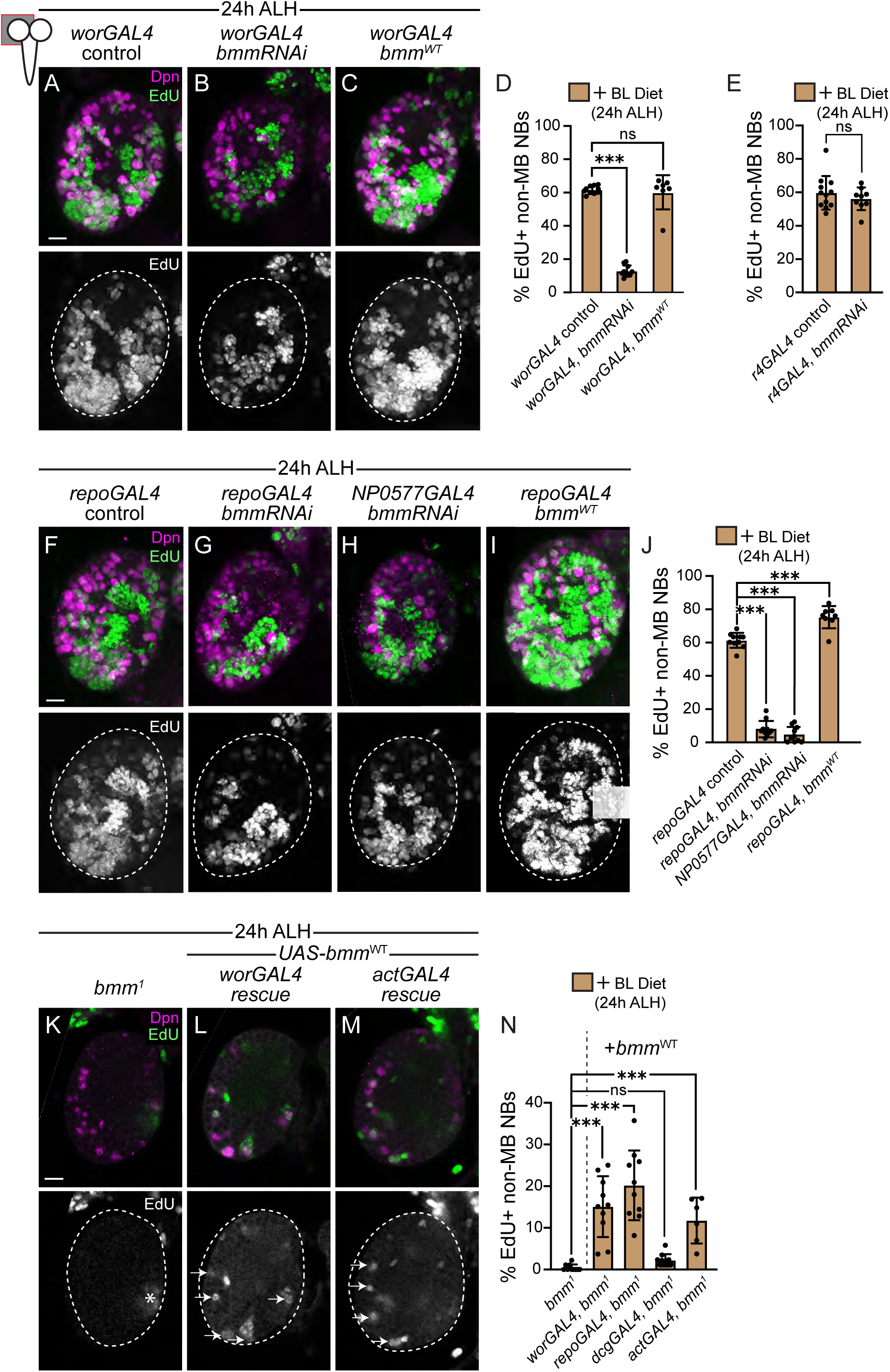
Lipid droplets in the freshly hatched larval brain are consumed locally to support neuroblast reactivation. (A-C,F-I,) MIPs of single brain hemispheres. Time points and genotypes listed above and markers within panels. Top, colored overlay with grayscale image below with brain hemispheres outlined. (D,E,J) Percentage of non-MB NBs that are EdU positive after 24 hours of BL feeding. (K-M) Single optical Z-images along the DV axis at similar locations after 24 hours of EdU feeding on BL-diet with quantification (N). Top panels, colored overlays with grayscale images below. (K) Asterisk indicates EdU positive neurons generated from MB NBs and arrows (L,M) indicate EdU positive reactivated non-MB NBs. Each data point (D,E,J,N) represents one brain hemisphere from one animal. Mean ± SD; (D,J,N) one-way ANOVA with Dunnett’s post hoc test versus control (***p≤0.05), ns (no significance), (E) Student two-tailed t-test. Scale bar in (A,F,K) equals 10μm. Panel by panel genotypes listed in Supplemental Table 1.

Knockdown of *bmm* in either neuroblasts or glia increased LD number (SF3A-J). We also overexpressed wildtype *bmm* (*UAS-bmm^WT^*) in neuroblasts (*worGAL4*) and glia (*repoGAL4*) to test whether LD breakdown is sufficient for reactivation. Overexpression of *bmm* in glia or neuroblasts led to reductions in brain LD numbers at FH stages, but only glial overexpression led to increased neuroblast reactivation 24h later (Fig. 2A,C,D,F,I,J and SF3K-M). Together, these results indicate that lipid droplets are consumed locally in neuroblasts and glia, and that neuroblast reactivation requires LD utilization in both cell types, reinforcing the idea that reactivation depends on coordinated neuroblast-glia interactions^15,40^.

Next, we re-expressed *bmm* in a cell- and tissue-specific manner in *bmm* mutants. At 24 hours after BL feeding, re-expression of *bmm* in neuroblasts (*worGAL4*) or glia (*repoGAL4*), but not fat body (*dcgGAL4*) modestly increased neuroblast reactivation compared to *bmm* mutants alone (Fig. 2K,L,N). Next, we re-expressed *bmm* ubiquitously using *actGAL4* and again found only a modest increase (Fig. 2M,N). In *bmm* mutants, LDs accumulate in both neuroblasts and glia (SF4A-E). Re-expression of *bmm* reduced LD number in a cell type specific manner (SF4F-J). Consistent with this, Bmm:eGFP was associated with LDs in glial rescue animals (SF4O, note: Bmm:eGFP LD association was not assayed in NB rescue animals). Ubiquitous expression reduced LD number at both FH stages and after 24 hours of BL feeding. We conclude that Bmm activity is tightly regulated, and that ubiquitous early expression likely depletes LDs prematurely, limiting rescue of neuroblast reactivation.

## Maternal lipid input controls neuroblast reactivation during larval feeding stages

Next, we asked whether brain LD number and neuroblast reactivation are maternally controlled. Both *Jabba* and *Lsd-2* maternal and maternal-zygotic mutants showed reduced LD number and neuroblast reactivation compared to OR animals, whereas *bmm* maternal-zygotic mutants died during embryogenesis (SF5A-C). Neuroblast reactivation is therefore a maternal effect phenotype, driven by changes in lipid droplet allocation and abundance in the egg. We next asked whether neuroblast reactivation could be modulated by maternal diet. We fed adult females a control BL-diet or a Lipid-depleted diet to alter egg lipid levels (Fig. 3A)^41,42^. After seven days on the Lipid-depleted diet, TAG levels relative to total protein were reduced by ∼20% in adult females (Fig. 3B). Progeny from these females also showed reduced LD number, both in early embryos and in the brain at FH stages (Fig. 3C-E and SF5D-F). Freshly hatched larvae from Lipid-depleted mothers reared on the BL-diet for 24 hours showed reduced neuroblast reactivation compared to those from BL-fed mothers (Fig. 3F-H). Thus, maternal diet influences stem cell behavior in offspring by controlling lipid availability for neuroblast reactivation.

**Figure 3:**
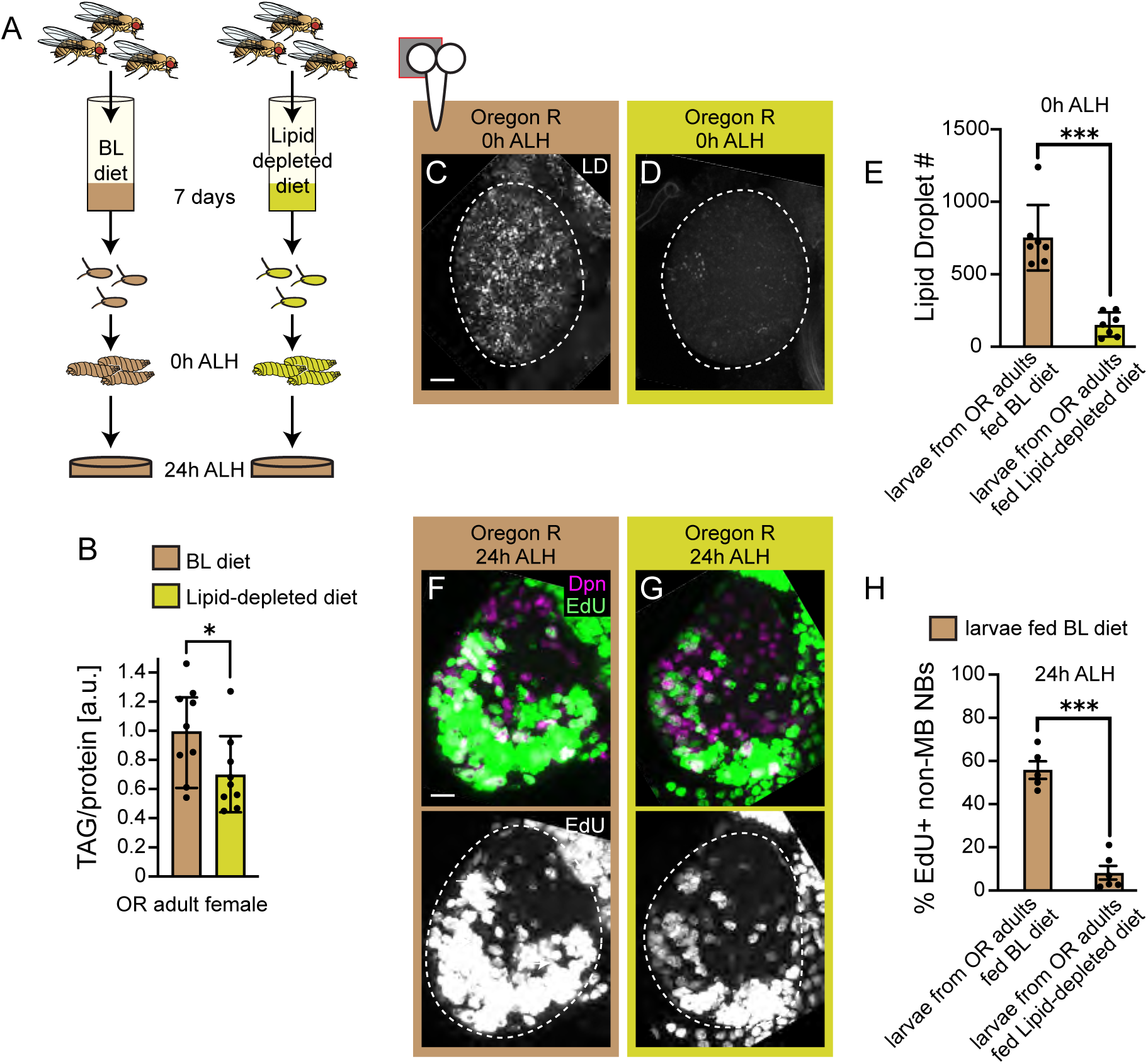
Maternal diet controls neuroblast reactivation during larval feeding stages. (A) Schematic of feeding paradigm. One day old Oregon R adults were transferred to and maintained on designated diets for seven days. Freshly hatched larvae from adults fed different diets were placed on the same BL-diet for 24 hours. (B) Normalized TAG levels in single adult females after seven days on designated diets. Each data point is one female. [a.u], arbitrary unit. (C,D) MIPs of single brain hemispheres of freshly hatched larvae (0h ALH) collected from adults fed different diets. LD image alone with brain hemispheres outlined. (E) Quantification of LD number in brain hemispheres. Each data point represents one brain hemisphere from one animal. (F,G) MIPs of single brain hemispheres after EdU feeding on BL food for 24 hours. Top panels, colored overlays with grayscale images below with brain hemispheres outlined. (H) Quantification of EdU positive neuroblasts after feeding on a BL-diet. Each data point represents one brain hemisphere from one animal. (B,E,H) Mean and S.D. Student two-tailed t-test (***p≤0.001). Scale bar (C,F) equals 10μm. Panel by panel genotypes listed in Supplemental Table 1.

## LD consumption fuels signaling and metabolism required for neuroblast reactivation

Consumption of LDs could supply lipids for membrane biogenesis, generate lipid-derived intermediates for signaling, or provide substrates for mitochondrial β-oxidation. During early larval stages, neuroblasts enlarge prior to S-phase re-entry and cortex glia undergo a concomitant expansion in surface area^12,13,15,38–40^. After 24 hours of feeding on a BL-diet, *Jabba*, *Lsd-2* and *bmm* mutant neuroblasts remained small compared to OR neuroblasts, similar in size to quiescent neuroblasts (SF6A). Cortex glial membrane surface area was also reduced, suggesting that LDs support membrane biogenesis (SF6B-E). To determine whether growth signaling was affected, we assayed components of the Insulin/PI3-kinase pathway. In both OR animals and *bmm* mutants, Dilp-2 (*Drosophila* insulin-like peptide) was synthesized and secreted from the IPCs, accumulating in the Dilp-2 recruiting neurons (DRNs) in each of the brain hemispheres (Fig. 4A-C). Dilp-2 is required for neuroblast reactivation and cortex glia growth and is released from the IPCs (insulin producing cells) both systemically and locally into the brain^13,15,18,43^.

**Figure 4:**
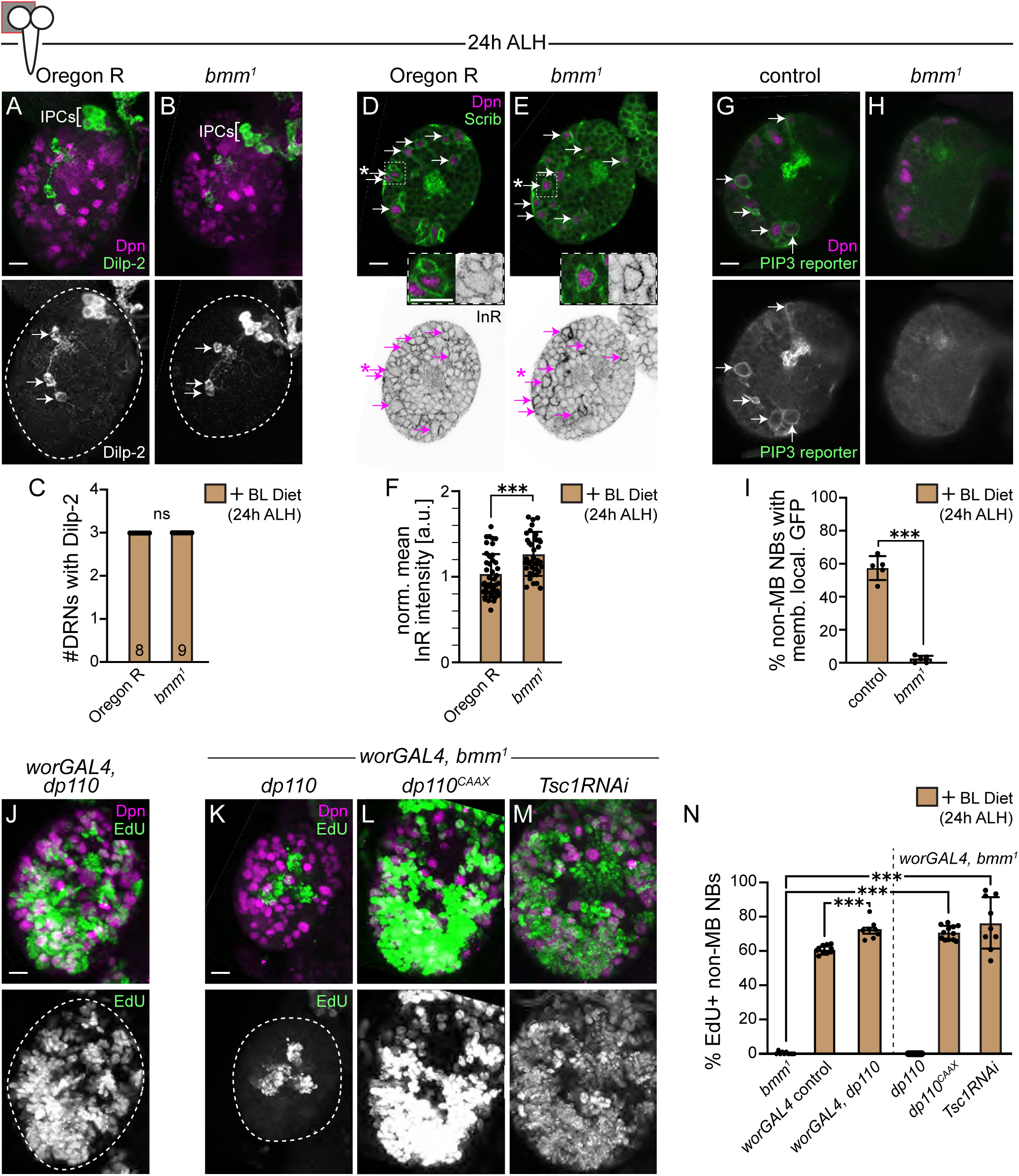
Maternally deposited lipid droplets promote early neural tissue growth and reception of Dilp nutrient signals during feeding stages. (A,B) MIPs of single brain hemispheres. Time points and genotypes listed above and markers within panels. Top, colored overlay with grayscale image below with brain hemispheres outlined. Arrows point to the DRNs (Dilp-2 recruiting neurons) which capture Dilps released from IPCs with quantification (C, n of brain hemispheres scored indicated in column). (D,E) Single Z images. Top, colored overlay and below same Z with inverted grayscale image showing InR labeling. Arrows point to non-MB neuroblasts, and in boxed outline shown at higher magnification below. In *bmm* mutants, neuroblasts have higher mean InR fluorescence along their plasma membranes compared to OR animals (quantified in F). (G,H) Single Z images with colored overlay (top) and grayscale image below. Arrows point to neuroblasts with reconstituted Venus PIP3 reporter expression along the plasma membrane with quantification (I). (J-M) MIPs of single brain hemispheres after EdU feeding on BL food for 24 hours with quantification (N). Top panels, colored overlays with grayscale images below. Each data point (C,I,N) represents one brain hemisphere from one animal and (F) one neuroblast. Mean ± SD; (C,F,I) Student two-tailed t-test (***p≤0.001), ns (no significance), [a.u], arbitrary unit. (N) one-way ANOVA with Dunnett’s post hoc test versus control (***p≤0.05). Scale bar (A,D,G,J,K) equals 10μm. Panel by panel genotypes listed in Supplemental Table 1.

We next assayed InR (Insulin-like receptor) expression and found that neuroblasts in both *bmm* mutants and OR animals express InR, but that InR levels were consistently higher along the plasma membrane in *bmm* mutant neuroblasts (Fig. 4D-F). Expression of InR is positively regulated by the transcription factor Foxo, which itself is negatively regulated by insulin/PI3-kinase signaling in flies and mammals^44^. Thus, induction of InR suggests that InR signaling is not active in *bmm* mutants, consistent with InR being a Foxo target gene.

To directly test PI3-kinase activity, we used a bi-molecular fluorescence complementation reporter that detects PIP_3_ at the plasma membrane. PI3-kinase is a lipid kinase and when active, phosphorylates PIP_2_ (phosphatidylinositol 4,5-bisphosphate) at the 3’ position to generate PIP_3_ (phosphatidylinositol 3,4,5-trisphosphate). PIP_3_ serves as a binding site for pleckstrin homology domain containing proteins, including Akt, whose recruitment to the plasma membrane leads to activation of TOR growth signaling and inhibition of Foxo-mediated transcriptional programs^45,46^. The bi-molecular fluorescence complementation reporter consists of two transgenes encoding N- and C-terminal halves of GFP fused to the pleckstrin homology domain of GRP and when PI3-kinase is active GFP is reconstituted along the plasma membrane where PIP_3_ is found^38,47,48^. After 24 hours of BL feeding, control neuroblast showed PIP3 sensor signal along the plasma membrane (Fig. 4G,I). In contrast, no PIP_3_ sensor signal was detected in *bmm* mutants (Fig. 4H,I). This indicates that PI3-kinase signaling is inactive in *bmm* mutant neuroblasts despite feeding.

We next tested whether forced activation of PI3-kinase could restore reactivation in *bmm* mutants. Expression of the catalytic subunit (*UAS-dp110*) in control neuroblasts increased reactivation after BL feeding, but *bmm* mutant neuroblasts failed to respond (Fig. 4J,K,N). In contrast, expression of membrane-targeted PI3-kinase, *dp110^CAAX^*, which anchors the catalytic subunit to the plasma membrane to continuously convert PI(4,5)P_2_ to PI(3,4,5)P_3_, independent of InR or lipid-dependent recruitment, restored reactivation (Fig. 4L,N and SF6F). Similarly, expression of *TSC1RNAi* which activates TOR independently of membrane localized PI3-kinase also restored neuroblast reactivation (Fig. 4M,N and SF6G). Together, these results suggest that strong membrane-localized PI3-kinase or downstream activation of TOR kinase is sufficient to bypass the requirement for LD breakdown, and that Bmm may function to support some aspect of the membrane environment required for PI3-kinase signaling.

Finally, to test whether LD breakdown supports mitochondrial metabolism, we examined *withered* (*whd*) mutants. *whd* encodes carnitine palmitoyltransferase I (CPTI), an enzyme in the carnitine shuttle that transfers long-chain fatty acids onto carnitine for transport into the mitochondrial matrix, where β-oxidation occurs^49^. Without CPT activity, fatty acids cannot enter mitochondria and are unavailable for oxidation. Like *bmm* mutants, *whd* mutants failed to reactivate neuroblasts after 24 hours of BL feeding, suggesting that LD breakdown contributes to reactivation by providing substrates for mitochondrial ATP production (SF6H-J).

## Glial Lsd-2 controls neuroblast reactivation non-autonomously through LD mobilization

Having established that LD breakdown is required locally in the brain and fuels both growth signaling and metabolism, we next investigated how LD mobilization is regulated at the cellular level. Using existing publicly available datasets, we identified genes that were differentially expressed between neuroblasts and glia in the L1 larval brain (SF7A)^50,51^. We cross-matched this list with gene ontogeny terms that are associated with LD storage, metabolism, and organization (Supplemental Table 2). We identified 29 genes, including Lsd-2 (vertebrate *perilipin-2*) that was expressed at 6-fold higher levels in glia compared to neuroblasts (SF7A). Lsd-2 encodes a LD surface protein that shields droplets from lipolysis by lipases such as Bmm^32,33,52^. Consistent with this role, Lsd-2 was associated with brain LDs in glia at both FH stages and 24 h after BL feeding, with increased LD-associated Lsd-2 in *bmm* mutants likely due to LD accumulation (Fig. 5A–D and SF7B). Lsd-2 was not associated with LDs in neuroblasts in control or *bmm* mutants (SF7C,D).

**Figure 5:**
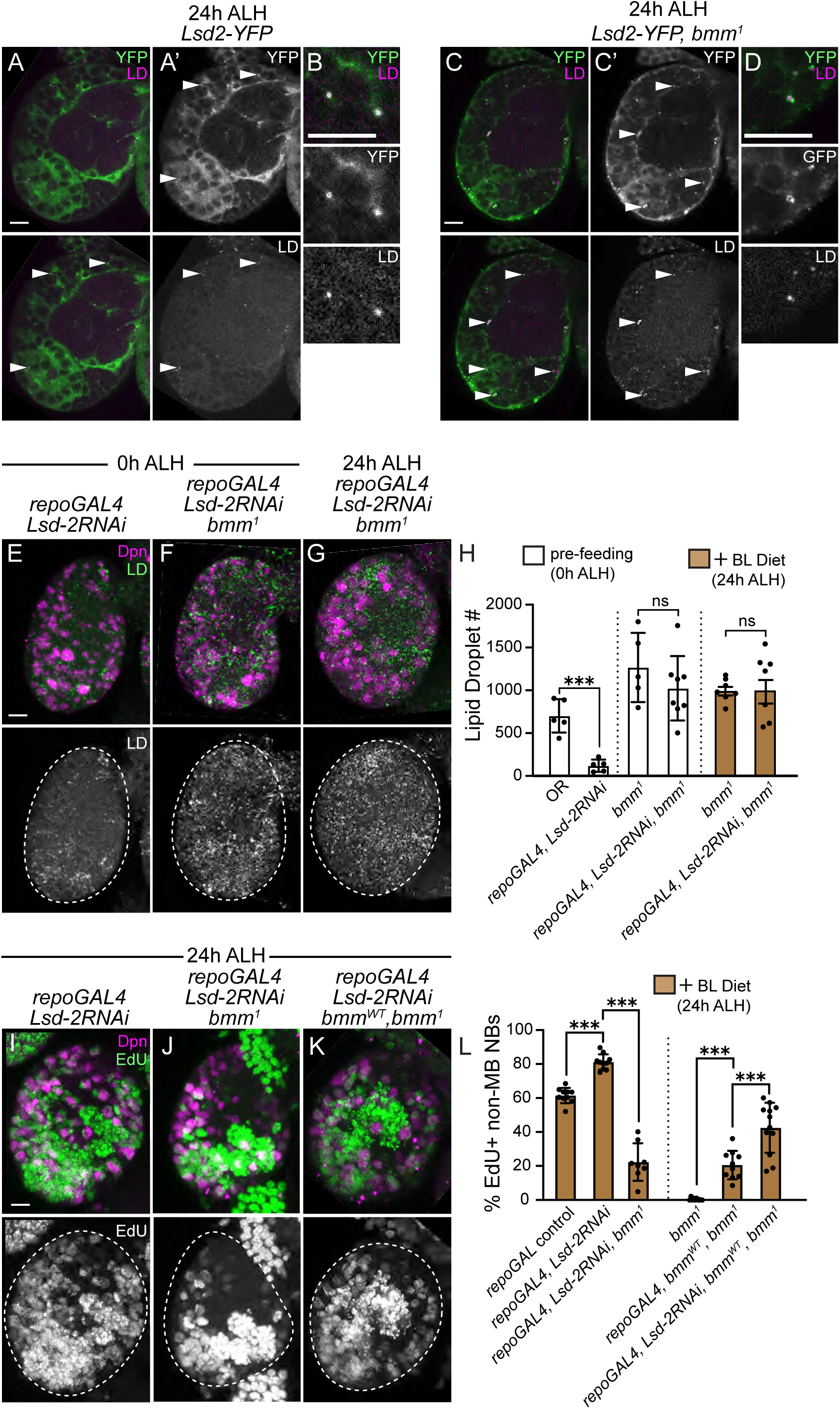
Lsd-2 in glia promotes LD breakdown and neuroblast reactivation. (A-D) Single Z images of brain hemispheres with higher mag LD images (B,D). Colored overlay, with grayscale images below, arrowheads mark some of the LDs associated with Lsd-2. Time points and genotypes listed above and markers within panels. (E-G) MIPs of single brain hemispheres. Top panels, colored overlays with grayscale images below with brain hemispheres outlined. (H) Quantification of LD number. (I-K) MIPs of single brain hemispheres after 24 hours of EdU feeding on BL-diet with quantification in (L). Each data point (H,L) represents one brain hemisphere from one animal. Mean ± SD; (H,L) one-way ANOVA with Dunnett’s post hoc test versus control (***p≤0.05), ns (no significance). Scale bar (A-D,E,I) equals 10μm. Panel by panel genotypes listed in Supplemental Table 1.

To test function, we reduced *Lsd-2* specifically in glia. Glial knockdown of *Lsd-2* (*UAS-Lsd-2RNAi*) reduced brain LD number and increased neuroblast reactivation after BL feeding, whereas knockdown in neuroblasts had no effect (Fig. 5E,H,I,L and SF7E). Increased reactivation was also observed in glial *Lsd-2* knockdown animals at earlier time points, 12h and 18h ALH (SF7F). When *Lsd-2* was knocked down in glia in a *bmm* mutant background, LD number and reactivation was similar to *bmm* mutants alone (Fig. 5F-G,J,L), indicating that Lsd-2 regulates LD mobilization upstream of Bmm. Finally, *Lsd-2* knockdown with *bmm* re-expression (*bmm^WT^*) in glia in *bmm* mutants modestly improved reactivation compared to *bmm* expression (*bmm^WT^*) alone (Fig. 5K,L). Together, these results indicate that Lsd-2 limits Bmm access to LDs in glia to regulate LD mobilization during NB reactivation, whereas LDs in neuroblasts are regulated by a distinct mechanism likely not involving Lsd-2.

### Discussion

In response to animal feeding, insulin-like peptides are synthesized and released, locally and systemically, leading to InR/PI3-kinase pathway activation and organism-wide increases in cell number and tissue size^1,14,16,18,43,53^. In the brain, these systemic and local cues drive quiescent neuroblasts back into the cell cycle to continue generating neurons needed for larval and adult neural function^12,13,15,38,40^. Beyond acquisition of nutrients through feeding, we show here that maternal lipids deposited into the egg and stored in LDs are also required to reactivate quiescent neuroblasts. We propose that both stored (maternal) and acquired (larval feeding) nutrients are required for neuroblasts to efficiently reactivate from developmental quiescence (SF8). If either nutrient source is disrupted or limited, neuroblasts still reactivate, but reactivation is delayed^12,13,15,38,40^. Whether delayed reactivation leads to reductions in numbers or types of neurons in later larval or adult brains remains an open question.

Lipid droplets contain a triacylglycerol (TAG)-rich core together with other neutral lipids, including cholesteryl esters, and are enclosed by a phospholipid monolayer that recruits lipid-metabolizing and regulatory proteins^23–25,54^. We find that both Bmm and Lsd-2 localize to the cytosolic phospholipid monolayer of LDs. Bmm selectively hydrolyzes the TAG component, releasing free fatty acids and glycerol, but does not act on cholesteryl esters or the structural monolayer^37^. Thus, the progressive reduction in LD number indicates activity of additional lipid catabolic pathways, including cholesteryl-ester hydrolases and LD-directed autophagy (lipophagy), to remove residual neutral lipids and remaining LD components. The free fatty acids released by Bmm activity could provide the carbon backbone needed for synthesizing new lipids, including phospholipids for membrane growth. Quiescent neuroblasts increase in size prior to cell-cycle re-entry, while glia expand their membrane surface area^12,13,15,38,40^. Free fatty acids from LD breakdown could also enter the glycerolipid pathway to generate diacylglycerol, phosphatidic acid, and phosphatidylinositol, which help organize membrane signaling microdomains and create phosphoinositide-rich regions that recruit and activate PI3-kinase^45,46^. When maternal LDs are reduced or when Bmm is knocked down, the neuroblast membrane environment might be insufficient to support PI3-kinase activation, consistent with our finding that membrane-targeted, catalytically active PI3-kinase restores signaling, whereas the cytosolic catalytically active form does not. At the same time, we show that LD-derived fatty acids fuel mitochondrial β-oxidation, as indicated by the failure of *whd* mutants to reactivate. Whether this requirement reflects a cell-autonomous need in neuroblasts or a non-autonomous role in surrounding glia or peripheral tissues remains unresolved. Together, these findings support a model in which lipid droplet breakdown provides the structural lipids needed for membrane biogenesis, the signaling lipids required for PI3-kinase/TOR pathway activation, and the metabolic substrates that fuel mitochondrial β-oxidation during neuroblast reactivation.

Reactivation from quiescence is regulated both cell-autonomously within neuroblasts and non-autonomously by surrounding glia^12,13,15,38,40^. PI3-kinase activation and cellular growth are required in both cell types, and LD breakdown is essential in each, yet only glial-specific LD breakdown is sufficient to initiate early neuroblast reactivation. This difference may reflect cell-type-specific modes of lipid storage or lipolytic regulation. Lsd-2 is detected on glial LDs but not on neuroblast LDs, and glial-specific knockdown of Lsd-2 accelerates and enhances neuroblast reactivation, whereas neuroblast-specific knockdown has no effect. These findings indicate that regulated LD turnover in glia modulates the timing of neuroblast reactivation and point to distinct lipid mobilization programs in glia and neuroblasts. Similar roles for Lsd-2 have been reported in other *Drosophila* tissues, including the fat body, oenocytes, and cortex glia, where LD-associated Lsd-2 modulates Hedgehog signaling to influence neural stem cell proliferation^27,55^. In mammalian systems, PLIN2, the Lsd-2 homolog, similarly limits LD turnover; in embryonic stem cells, PLIN2 degradation upon exit from pluripotency promotes lipid hydrolysis, alters mitochondrial lipid composition, and reduces acetyl-CoA and histone acetylation, thereby triggering differentiation^56^. Together, these observations suggest that controlled LD mobilization through Lsd-2/PLIN2 may represent a conserved mechanism linking lipid metabolism to developmental transitions.

A key finding is that maternal lipid input establishes the reservoir of LDs available to the larval brain, and maternal diet modulates this reservoir, reinforcing the concept that maternal physiology and nutritional state influence offspring development. Dietary fatty acids strongly influence phospholipid composition in membranes throughout the animal, providing a mechanistic basis for how maternal provisioning shapes early developmental environments^41^. Similar principles operate in other systems in which stored lipids sustain survival and govern developmental timing, including mammalian diapause and nematode dauer^6,20,31,57,58^. In diapause mouse embryos and in quiescent mammalian neural stem cells, stored lipids serve as a primary energy source^59–61^. Although it remains unclear whether free fatty acids or LD abundance act as upstream triggers for quiescence, free fatty acids are required for stem cell maintenance and for reactivation from quiescence and diapause across a range of species. Yeast exiting stationary phase likewise consume LDs rapidly to support renewed proliferation and increased cell number^62^. What controls LD breakdown in quiescent stem cells, including *Drosophila* neuroblasts, is not yet known. However, in adult rodents, both exercise and caloric restriction promote LD mobilization and are associated with enhanced neurogenesis, suggesting that metabolic cues regulating LD turnover may act broadly to couple physiological state with stem cell activity^63–66^.

Together, these findings place lipid droplets at the center of the developmental program that governs neuroblast reactivation, revealing that maternal provisioning, glial lipid mobilization, and mitochondrial fatty acid use collectively couple organismal physiology to stem cell behavior in the early brain. Future work will be needed to define which lipid species are most critical, how LD allocation across brain cell types is established, and whether manipulating maternal diet or glial lipid metabolism can be used to influence stem cell behavior across development and in other organisms.

## Acknowledgements

We are grateful to the following people and centers for providing reagents and resources. We thank Christoph Heier (Institute of Molecular Biosciences, University of Graz, Austria), Michael Welte (Department of Biology, University of Rochester, New York), the Bloomington *Drosophila* Stock center (BDSC), and the Kyoto Drosophila Stock center for providing fly strains. We thank Chris Doe, Eric Rulifson, and the Developmental Studies Hybridoma Bank (DSHB) for providing antibodies. We thank Karsten Siller and UVA research computing for customized Fiji plugins. We thank Tony Spano for help with diets. This work was supported by the NIH (S.E.S.-R35GM141886, M.L.B.-R01DK123433, K.E.D.-T32DK007646) and the Jefferson Scholars Foundation (K.E.D.).

## Materials and Methods

### Drosophila strains, genetics, staging, and diet

Animals were raised in uncrowded conditions at 25×C on Bloomington (BL) food in controlled light dark cycles (12 hours on/12 hours off) unless stated otherwise. The BL-diet used was commercially purchased from Archon Scientific and is referenced as the W1 recipe. The following strains were used: *Oregon R* (BDSC #5)*, bmm*^1^(^37^), *Df(3L)ED217* (BDSC #8074), *jabba^DL^* (^36^), *Lsd2^KG^*^00149^ (BDSC #13382), *klar^YG^*^3^ (^34^), *UAS-Lsd2RNAi* (BDSC #32846), *UAS-bmmRNAi* (BDSC #25926), UAS-*dp110* (BDSC #25914), *UAS-dp110CAAX* (^67^), *UAS-TscRNAi* (BDSC #52931), *UAS-bmm WT* (BDSC #76600), *UAS-N-Venus-PH-GRP(*^48^), *UAS-C-Venus-PH-GRP(*^48^), *UAS-mCD8GFP* (BDSC #5137), *worGAL4(*^68^), *NP0577GAL4* (Kyoto #112228), *repoGAL4* (BDSC #7415), *r4GAL4(*^69^), *dcgGAL4* (BDSC #7011), *actin5CGAL4* (BDSC #4414), *Lsd2-YFP* (Kyoto #115301), *bmm-GFP* (BDSC #94600), *UAS-bmm:eGFP* (BDSC #98110), and *whd*^1^ (BDSC #441). Mutant alleles and transgenes were maintained over *actin:GFP* balancers for genotyping progeny. Adults were housed in condos and eggs collected on yeasted grape juice agar plates. Freshly hatched larvae (0-2 hours ALH) were collected from plates and either dissected or transferred to meal caps containing BL food (Archon Scientific W1) supplemented with or without EdU. The Lipid-depleted diet consisted of 10% yeast autolysate, 10% glucose, and 10% agar^41,42^.

### Immunofluorescence and confocal imaging

Larval brains were dissected and fixed in 4% EM grade formaldehyde in PEM (Pipes, EGTA, Magnesium chloride) buffer with 0.1% Triton-X for 20 minutes as described previously^15,38,39^. Tissues were washed in PBT (1X PBS with 0.1% TritonX-100) and blocked overnight at 4×C in PBT with 10% normal goat serum. Primary antibodies used in this study include: chicken anti-GFP (1:500, Abcam), rat anti-Deadpan (1:100, Abcam), rabbit anti-Scribble (1:1000, gift, used as a membrane maker), mouse anti-Repo (1:5, DSHB, 8D12), guinea-pig anti-InR (1:300), rabbit anti-Dilp2 (1:1000, gift), mouse anti-Dlg (DSHB, 1:40). Primary antibodies were detected using Alexa Fluor-conjugated secondary antibodies. LDs were detected using 1:200 dilution in PBS of Nile Red (stock 500mg/ml in acetone) after final secondary antibody wash for 40 minutes in the dark. For EdU labeling, animals were fed 0.1mg/ml EdU mixed in with BL food (or LDM food) and animals fed for designated amounts of time. EdU detection was performed for 30 minutes after final secondary antibody wash in the dark as described previously^15,38,39^. Brains were cleared in glycerol based antifade and imaged using a Leica SP8 laser scanning confocal microscope equipped with a 63X, 1.4 NA oil-immersion objective. For glia surface measurements, Z stacks were acquired at 0.5 μm steps using the same confocal settings. All other confocal data were collected at 1.0 μm steps. Images were processed using Fiji and figures assembled using Adobe Photoshop and Illustrator. LD staining in embryos was carried out in the absence of methanol as described previously^2,35^.

### Quantification of neuroblast EdU number, LD number, and glia membrane surface area

For neuroblast EdU quantification, the number of EdU-positive, Dpn-positive neuroblasts were counted using the “cell counter” plugin in Fiji and divided by the total number of Dpn-positive neuroblasts per brain hemisphere. Neuroblast diameter was calculated based on the average length of two perpendicular lines drawn through the center of the neuroblast at its widest point. We used custom Fiji plugins “Droplets” and “Glia Membrane” to quantify both LD number and glial membrane surface area. Both plugins are publicly available at https://github.com/neuroblastlab/lipid_droplets.git and https://github.com/neuroblastlab/glia_membrane.git. For EdU counts, all neuroblasts in the central brain region were assayed from the dorsal to ventral brain surface. For neuroblast diameter, LD number, and glia membrane surface area, only the top 50 μms from the dorsal surface were included in the quantification. LDs in neuropil glia were manually counted through individual Z-stacks. LDs were counted in neuropil glia because these cells are discrete and easily identifiable, whereas cortex glia are intermingled with neuroblasts and neurons, preventing unambiguous assignment of LDs to cortex glia at early stages. All LDs were counted in the neuropil and included in the quantification.

### Triglyceride and protein measurements

Single adult females were sonicated three times for ten seconds each time in 140 mM NaCl, 50 mM Tris-HCl, pH 7.4, 0.1% Triton X-100 with protease inhibitors (Roche). Following clearing by centrifugation at 4°C, supernatants were transferred to new tubes. Triglyceride (Liquicolor Test, Stanbio) and protein (BCA assay, Pierce) were measured in each sample, and triglyceride levels were normalized to protein levels.

### RNA sequence analysis

Differential gene expression (DGE) between glia and neuroblast cell types was determined using available single cell RNA seq data^50,51^. Cells within each biological replicate were grouped by cell type annotations (neuroblasts, dpn and glial, repo) and their raw expression counts aggregated using Seurat R package. This resulted in a single count matrix per cell type per biological replicate, which simulates a bulk RNA-seq experiment. Differential gene expression analysis was conducted using DESeq2 package in R. Genes with an adjusted p-value < 0.05 and a log2 fold change greater than 0.5 were considered significantly differentially expressed. DGE between glia and neuroblasts was visualized using a volcano plot which was generated using ggplo2 R package.

### Statistical analysis

Student’s t-tests and one-way ANOVAs with Dunnett’s post hoc test were performed using Prism 10. For box plots, the boundary of the box closest to zero indicates the 25th percentile, a line within the box marks the median, and the boundary of the box farthest from zero indicates the 75th percentile. Whiskers (error bars) above and below the box indicate the maximum and minimum, respectively. The data in plots and the text are presented as means ± SD. In general, only statistically significant differences are indicated in figures; all other comparisons were not significant which in some cases were designated NS.

## Supplementary Figure Legends

**Supplemental Figure 1:**
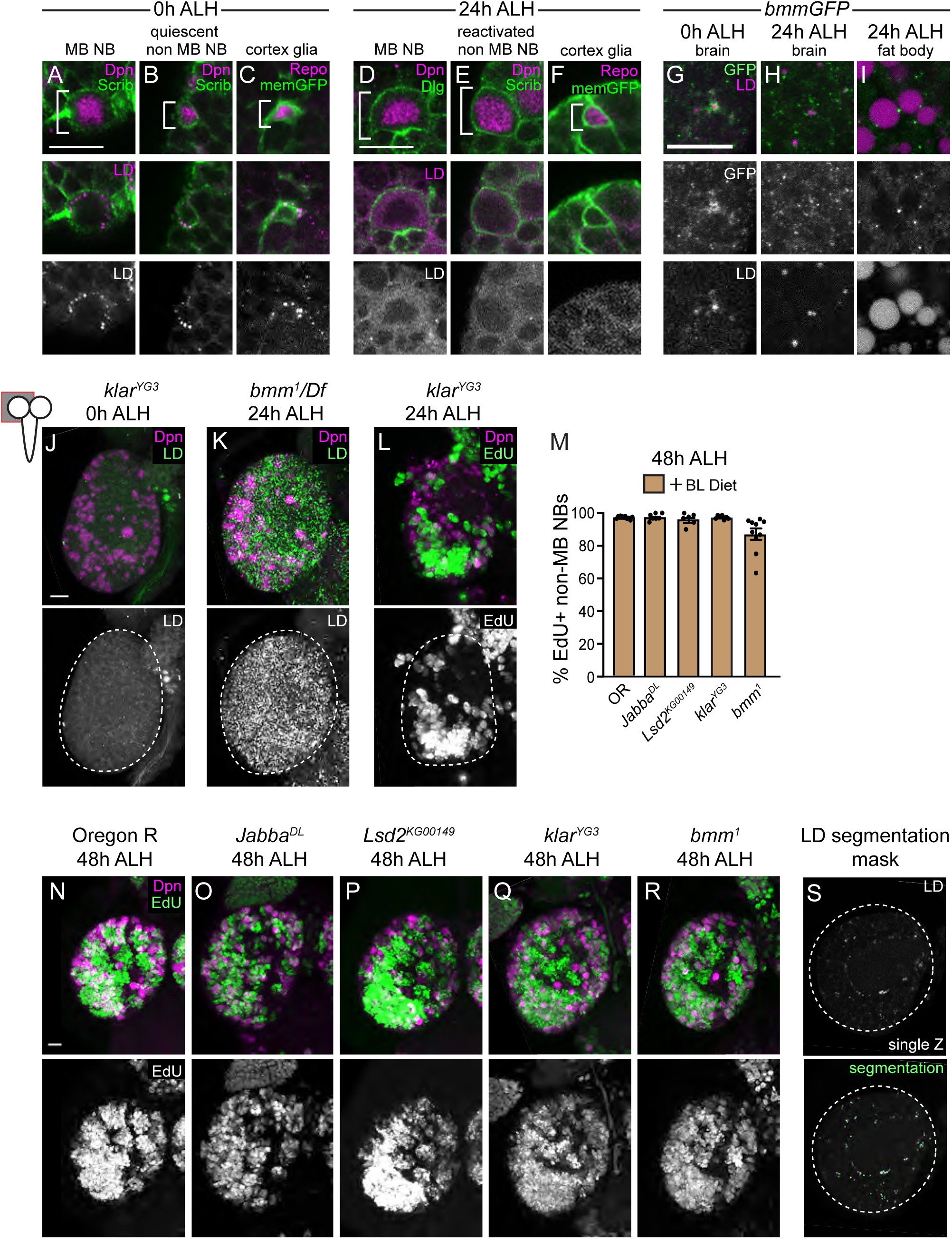
Related to Figure 1, Lipid droplet consumption correlates with neuroblast reactivation. (A-F) Single Z images of the distribution of LDs in neuroblasts (non-MB neuroblasts and MB neuroblasts) in OR animals and in glia expressing mCD8GFP. Cell type marked with white bracket. (G-I) LD-associated Bmm at 0 and 24h ALH. (J-L) MIPs of single brain hemispheres. Top panels, colored overlays with grayscale images below with brain hemispheres outlined. Time points and genotypes listed above and markers within panels. (M) Quantification of EdU percentage at 48 hours ALH in BL-fed animals in panel M-Q. Each data point represents one brain hemisphere from one animal. Mean and S.D. (N-R) MIPs of single brain hemispheres. (S) Example of LD segmentation in OR control animal used for quantification with single Z grayscale image on top with LD segmentation (green) overlay below. Scale bars in panels A,D,G,J,N equal 10 μm. Panel by panel genotypes listed in Supplemental Table 1.

**Supplemental Figure 2:**
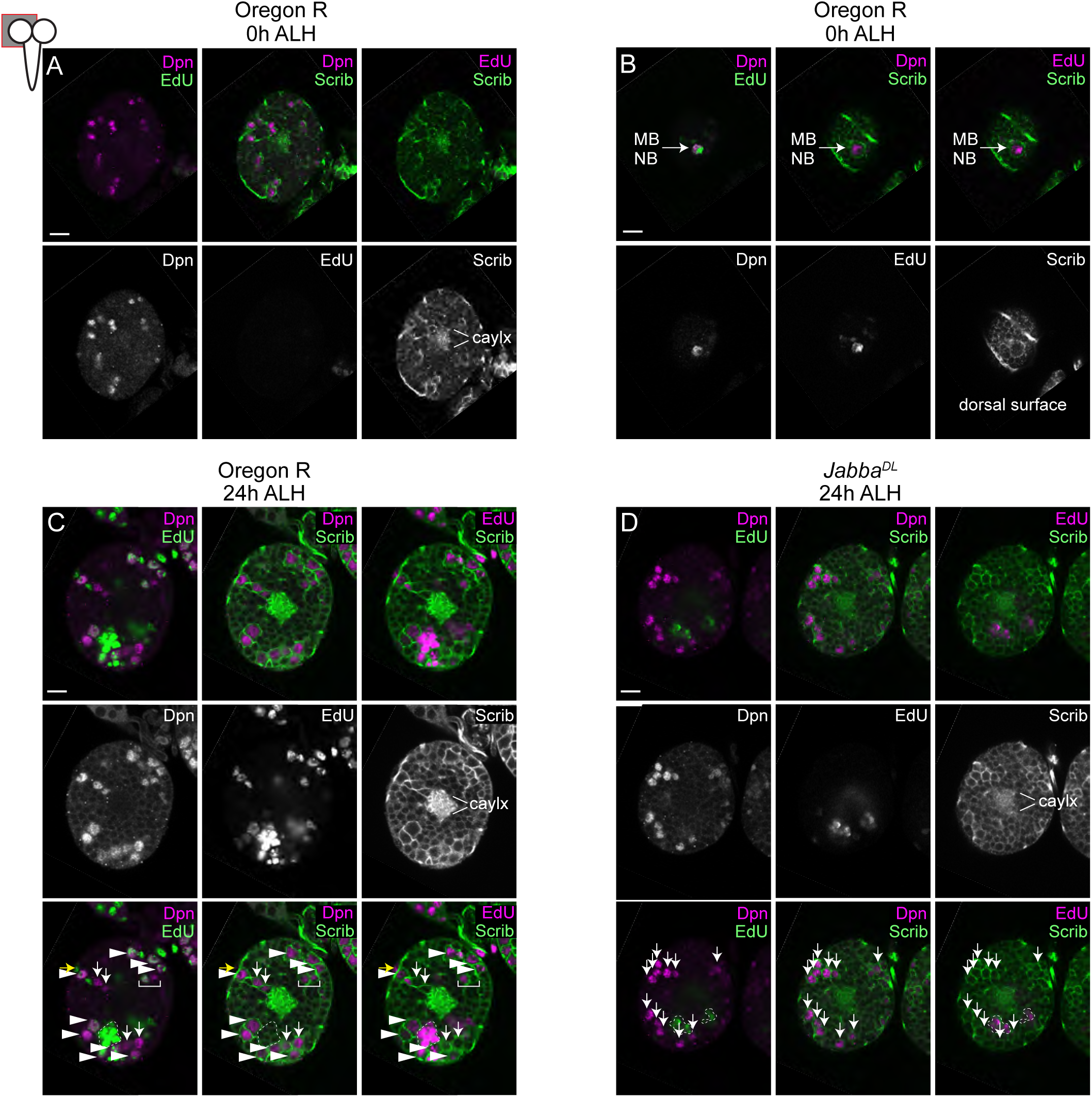
Related to Figure 1, Lipid droplet consumption correlates with neuroblast reactivation. (A-D) Single optical sections of EdU-labeled neuroblasts at indicated times and genotypes. Top row, colored overlay with single greyscale images below. In C and D, colored overlay is shown again with quiescent neuroblasts labeled with arrows and proliferating neuroblasts marked with arrowheads. EdU-labeled neuroblast neuron progeny are outlined in dashed white. White bracket indicates INPs of Type II lineage. Using the calyx as a landmark, A,C,D are similarly located, with B showing the dorsal brain surface with one of the four EdU-labeled MB neuroblasts. Scale bar in panels (A-D) equals 10 μm. Panel by panel genotypes listed in Supplemental Table 1.

**Supplemental Figure 3:**
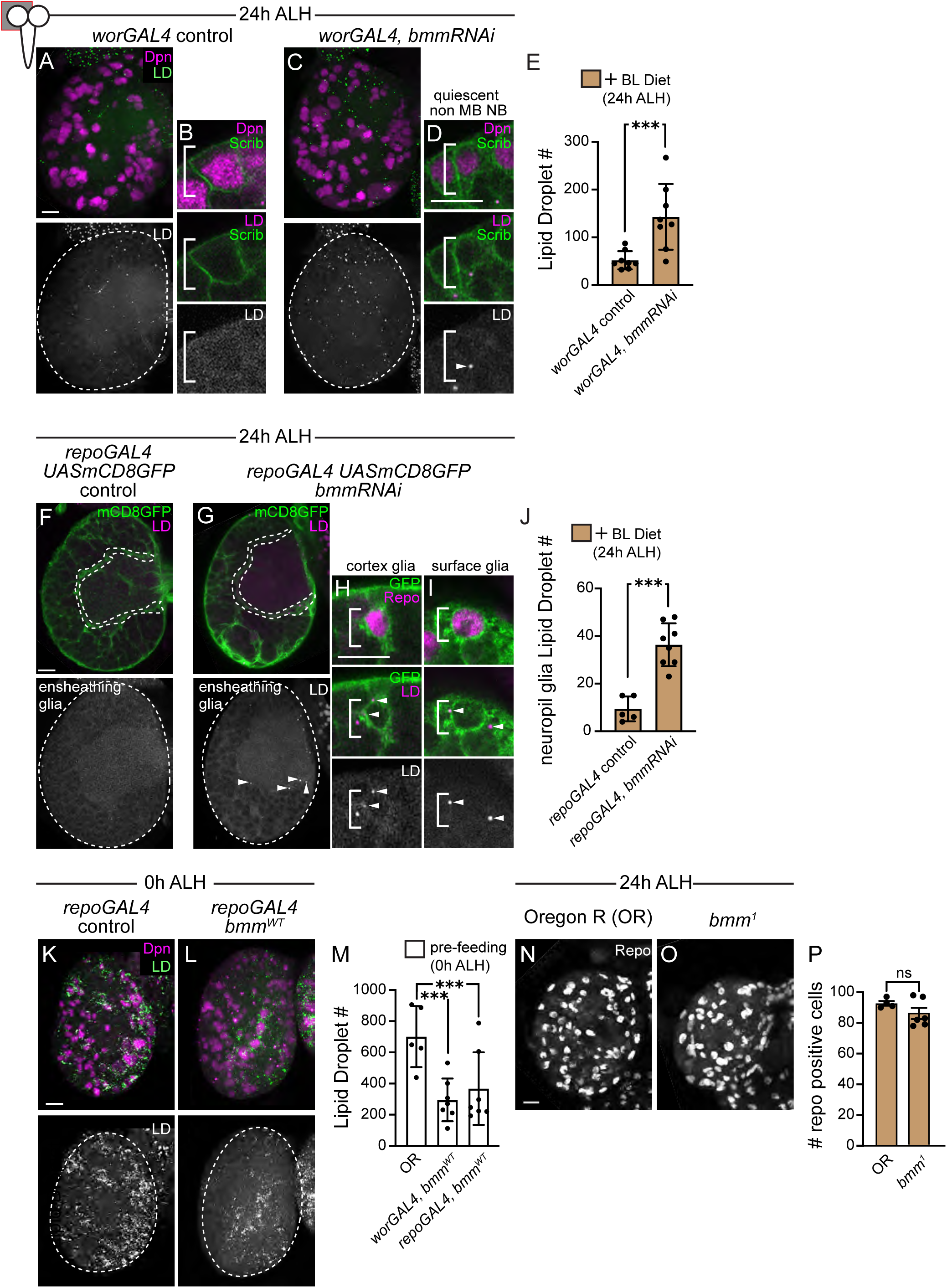
Related to figure 2, Lipid droplets in the freshly hatched larval brain are consumed locally to support neuroblast reactivation. (A,C) MIPs of single brain hemispheres in control and *bmmRNAi* animals with single Z images of neuroblasts at higher magnification (B,D). Top panels, colored overlays with grayscale images below with brain hemispheres outlined, neuroblasts in brackets, and LDs marked with arrowhead. (E) Quantification of LD number. (F,G) Single Z images of brain hemispheres in control and *bmmRNAi* animals with single Z images of glial types at higher magnification (H,I). Neuropil glia outlined in white (F,G) with quantification of LD number in this region in J. (K,L) MIPs of single brain hemispheres. Top panels, colored overlays with grayscale images below with brain hemispheres outlined. (M) Quantification of LD number. (N,O) MIPs of single brain hemispheres showing glia (Repo positive) in OR and *bmm*^1^ mutants with quantification of number of glia cells per brain hemisphere (P). Repo positive glia Each data point (E,J,M,P) represents one brain hemisphere from one animal. Mean ± SD; (E,J,P) Student two-tailed t-test (***p≤0.001); (M) one-way ANOVA with Dunnett’s post hoc test versus control (***p≤0.05). Scale bars are all 10 μm. Panel by panel genotypes listed in Supplemental Table 1.

**Supplemental Figure 4:**
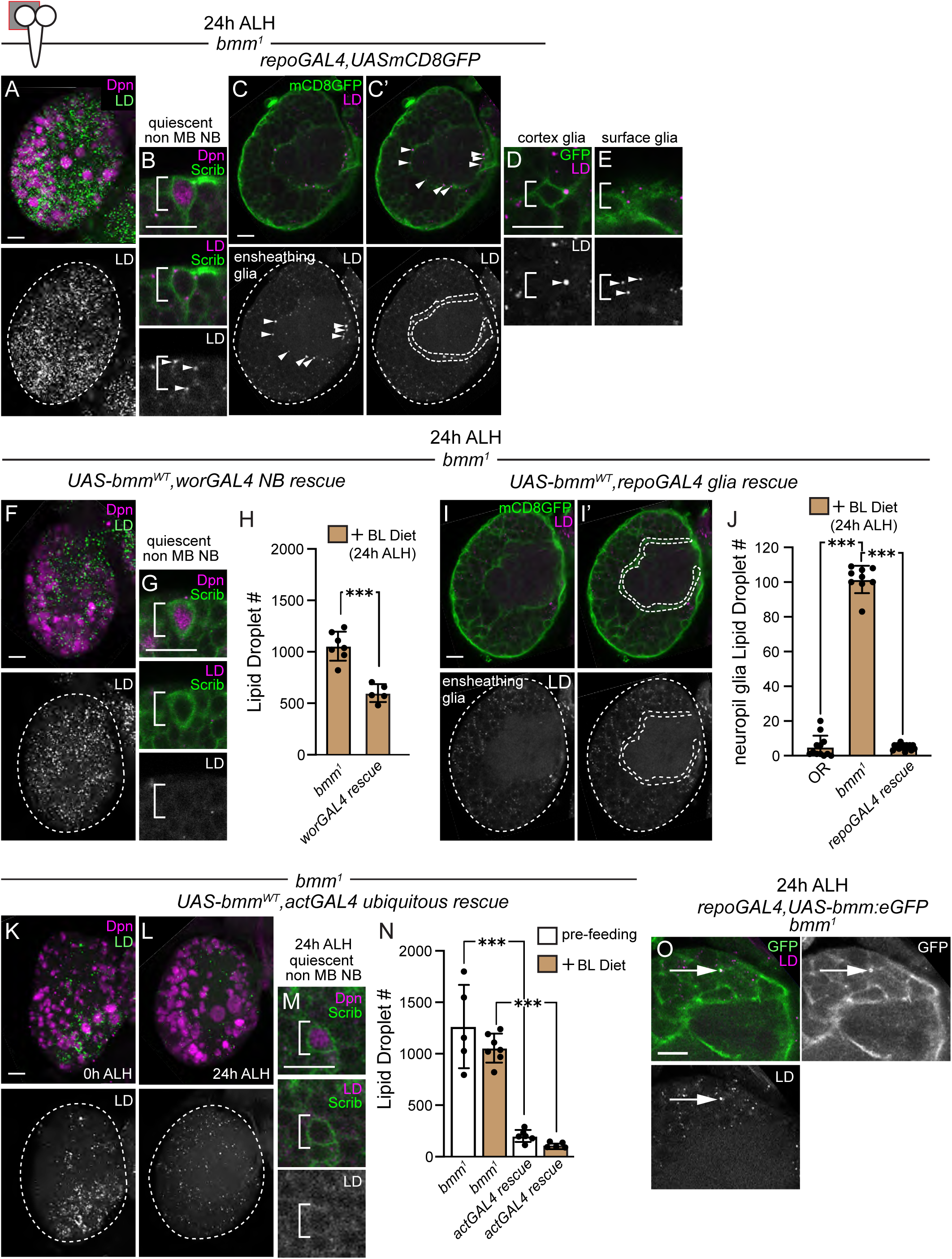
Related to figure 2, Lipid droplets in the freshly hatched larval brain are consumed locally to support neuroblast reactivation. (A) MIP and (C,C’) single Z images of a single brain hemisphere of homozygous *bmm*^1^ mutants showing LDs in some cell types (B,D,E and neuropil glia, white outline in C’). Top panels, colored overlays with grayscale images below with brain hemispheres outlined. Arrowheads indicate LDs and white brackets cell type. (F-N) Visualization and quantification of LDs in *bmm^WT^* rescue animals. Time points and genotypes listed above and markers within panels. All are cell type specific rescue in the *bmm*^1^ homozygous mutant background. Top panels colored overlay with grayscale images below and cell types at higher magnification to the right. (F-H) Neuroblast specific rescue (*worGAL4*), (I-J) glial specific rescue (*repoGAL4*), and (K-N) ubiquitous zygotic rescue (*actGAL4*). (O) High-magnification of single Z showing co-localization of *bmm:eGFP* with LDs in a *bmm*^1^ glial rescue animal. Each data point (H,J,N) represents one brain hemisphere from one animal. Mean and S.D. (H) Student two-tailed t-test (***p≤0.001); (J,N) one-way ANOVA with Dunnett’s post hoc test versus control (***p≤0.05). Scale bars are all 10μm. Panel by panel genotypes listed in Supplemental Table 1.

**Supplemental Figure 5:**
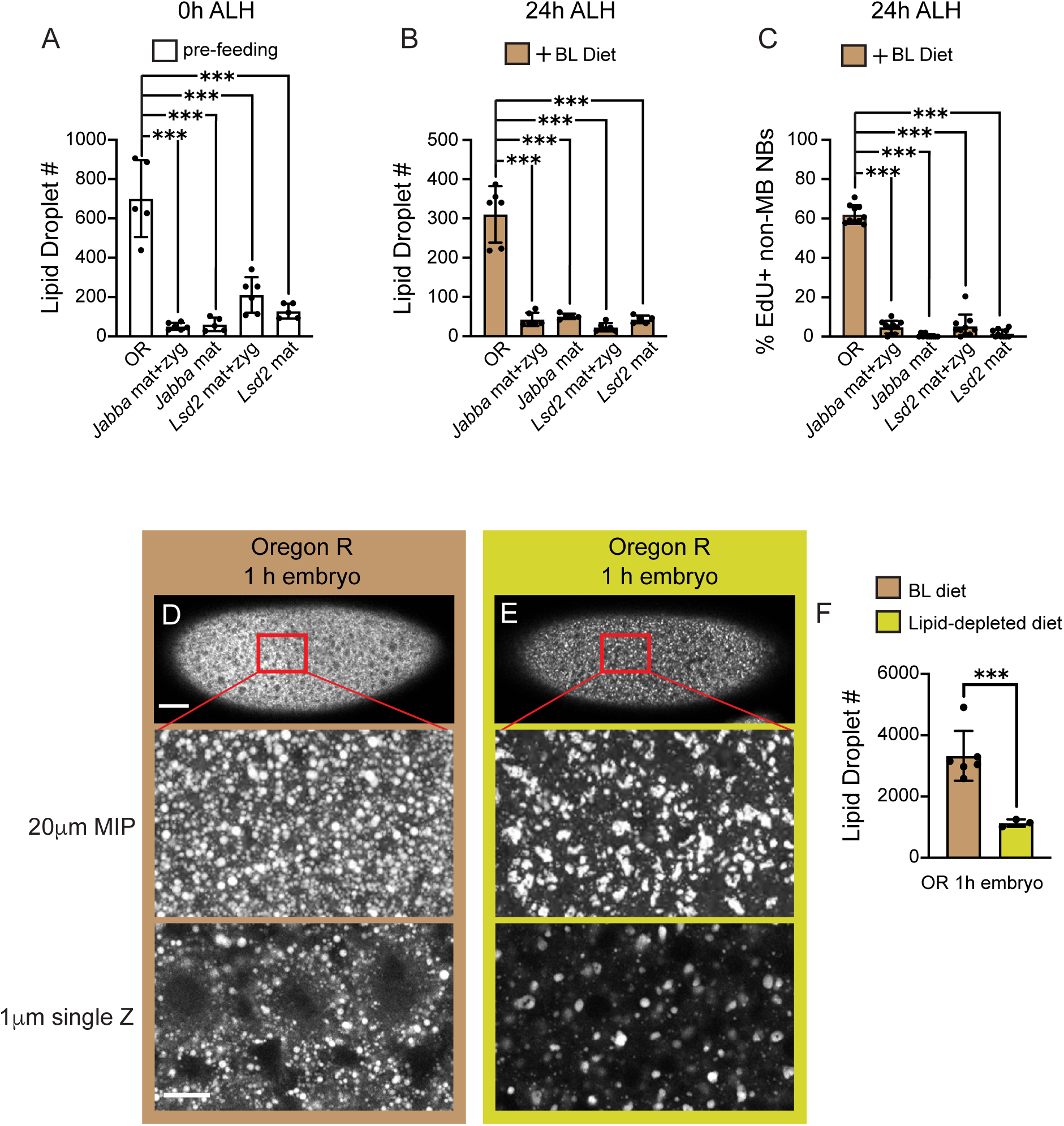
Related to figure 3, Maternal diets control neuroblast reactivation during larval feeding stages. (A,B) Quantification of lipid droplet number and percentage of EdU positive non-MB NBs (C) in maternal, zygotic mutants compared to maternal mutants. Genotypes listed below and time points above. Each data point represents one brain hemisphere from one animal. Mean and S.D. one-way ANOVA with Dunnett’s post hoc test versus control (***p≤0.05). (D-F) LD visualization in embryos of moms fed BL-diet versus Lipid-depleted diet with quantification. Each data point (F) is one embryo, mean and S.D., Student two-tailed t-test (***p≤0.001). LDs quantified from equal sized areas (125μm^2^). Scale bar (D) equals 50μm top panel for whole embryo and 10μm bottom panel for LD high magnification image. Panel by panel genotypes listed in Supplemental Table 1.

**Supplemental Figure 6:**
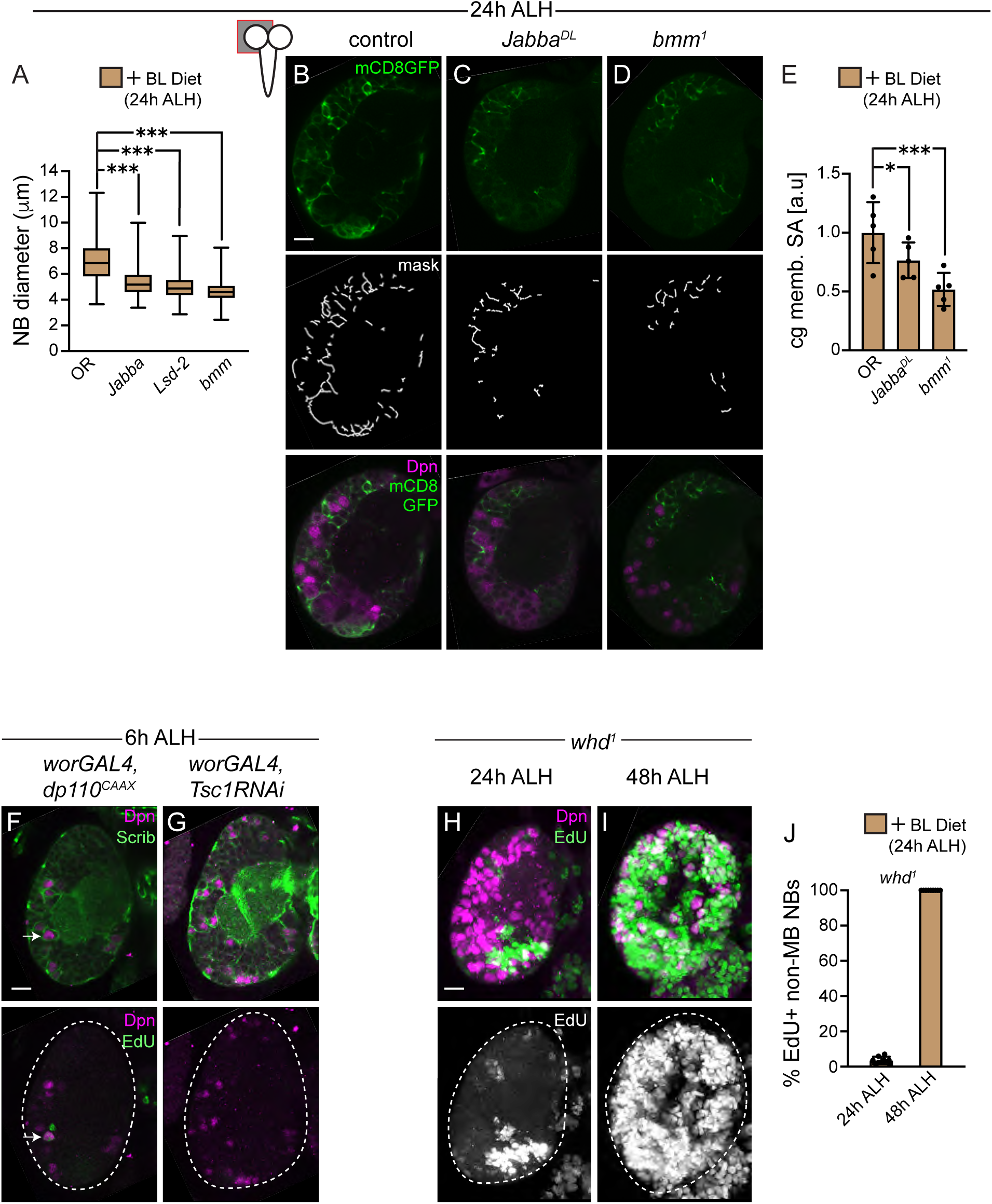
Related to figure 4, Maternally deposited lipid droplets promote early neural tissue growth and reception of Dilp nutrient signals during feeding stages. (A) Neuroblast size based on cell diameter. Each box and whiskers represent 150 neuroblasts quantified from three brain hemispheres from three animals (50 each). (B-D) Single optical sections of brain hemispheres. Top panels show membrane tagged GFP (mCD8GFP) driven by *NP0577GAL4* (cortex glia) and middle panels are masks used in quantification of cortex glia membrane surface area (E). (F,G) Single Z images along the DV axis at similar locations after 6 hours of EdU feeding on BL-diet. Time points and genotypes listed above and markers within panels. Arrowhead marks a prematurely dividing non-MB neuroblast. (H,I) MIPs of single brain hemispheres after EdU feeding on on BL food for 24 or 48 hours with quantification (J). Each data point (E,J) represents one brain hemisphere from one animal. Mean and S.D., one-way ANOVA with Dunnett’s post hoc test versus control (***p≤0.05). Scale bars are all 10μm. Panel by panel genotypes listed in Supplemental Table 1.

**Supplemental Figure 7:**
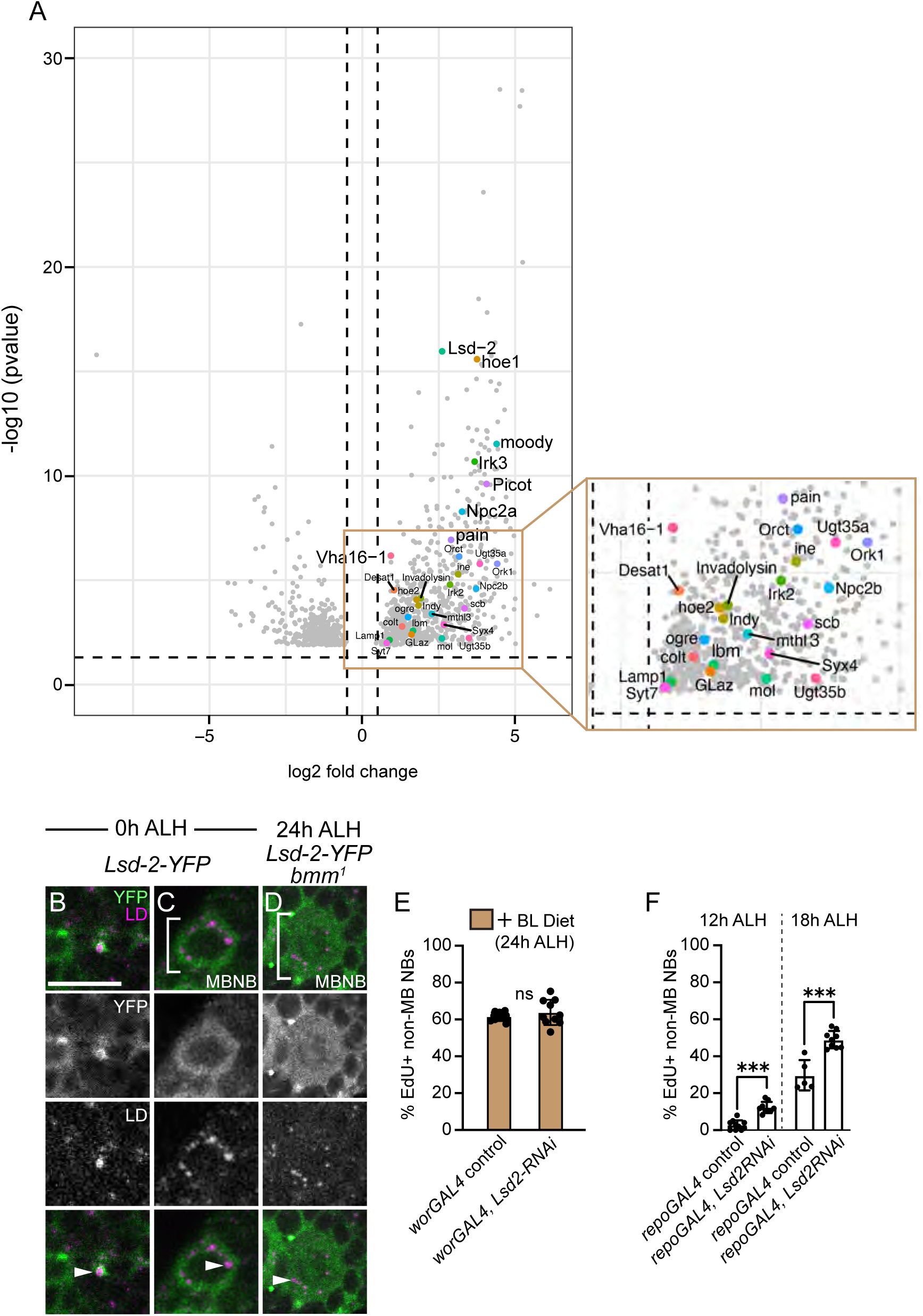
Related to figure 5, Lsd-2 in glia promotes LD breakdown and neuroblast reactivation. (A) Volcano plot showing differentially expressed genes (DEG) between glia and neuroblasts in first instar larvae. X-axis represents log2 fold change and Y-axis -log10 (p-value). Cut off for DEG was set at p-value <0.05 and log2 fold change > 0.5 (dashed lines). Labeled genes were only upregulated in glia and associated with GO terms describing lipid droplet metabolism. Inset was drawn for clarity. (B-D) High magnification images of LD-associated Lsd-2 in the brain. Time points and genotypes listed above and markers within panels. Top panels, colored overlays with grayscale images below. (E,F) Quantification of the percentage of EdU positive non-MB NBs after feeding on BL-diet for designated times. Each data point (E,F) represents one brain hemisphere from one animal. Mean and S.D. Student two-tailed t-test (***p≤0.001). Scale bar, B equals 10μm. Panel by panel genotypes listed in Supplemental Table 1.

**Supplemental Figure 8:**
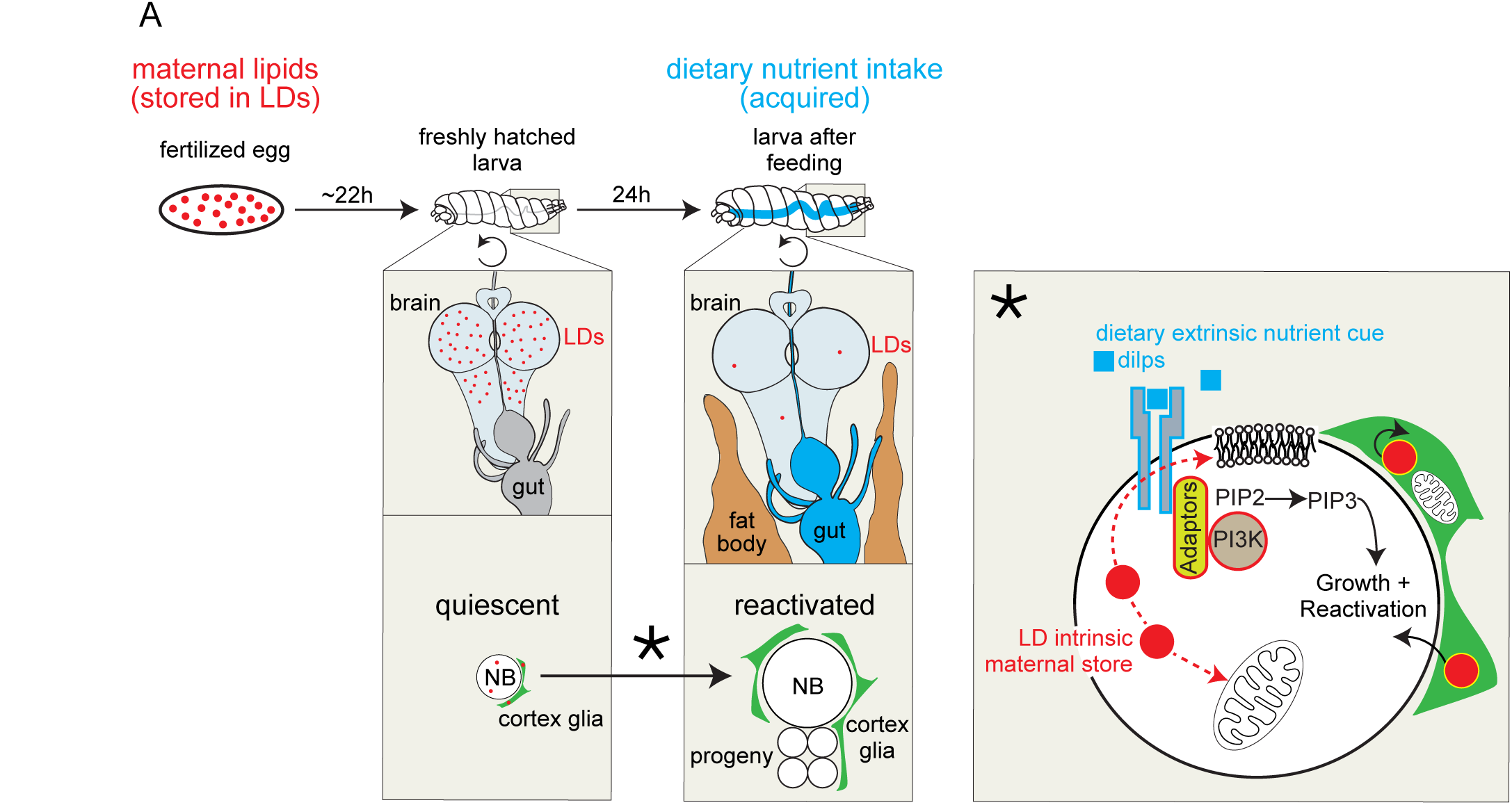
Related to figure 5, Lsd-2 in glia promotes LD breakdown and neuroblast reactivation. (A) Model figure. LDs (red spheres) are deposited into the egg. Maternally deposited lipids are consumed during early larval life to support neuroblast reactivation in conjunction with nutrients acquired through feeding. In freshly hatched larvae (before feeding, gut in grey), LDs are found throughout the brain in both neuroblasts and glia. At this time, neuroblasts are quiescent except for MB+VL neuroblasts and cortex glia (green) have little to no membrane elaboration. Twenty four hours later, after larval feeding (gut in blue), neuroblasts increase in size and resume proliferation, while cortex glia elaborate their membranes. During reactivation, LDs are broken down and utilized to support membrane growth, signaling, and metabolism. LD breakdown in both neuroblasts and glia supports neuroblast reactivation (see discussion). The dynamics of LD breakdown is differentiatedly regulated between glia and neuroblast cell types based on a requirement of Lsd-2 (yellow outline of LDs in glia) in glia but not neuroblasts. It remains unclear how LD breakdown in glia supports neuroblast reactivation (see discussion).

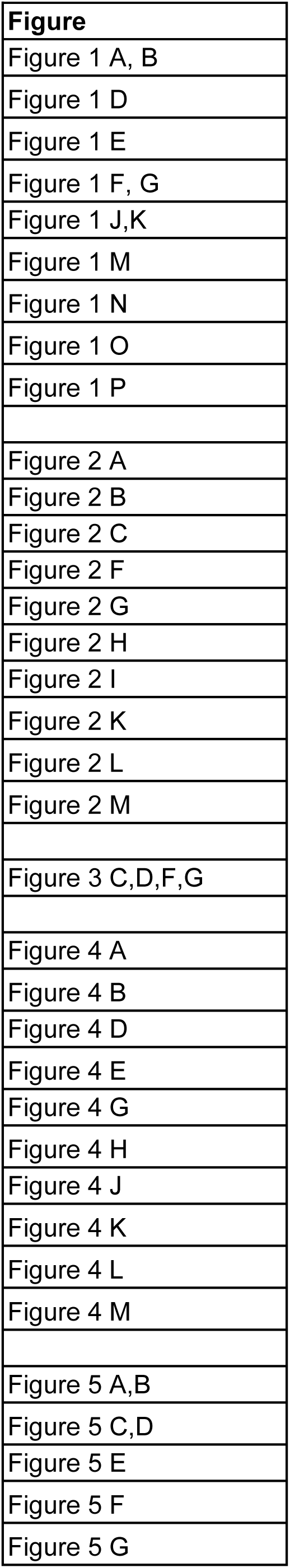
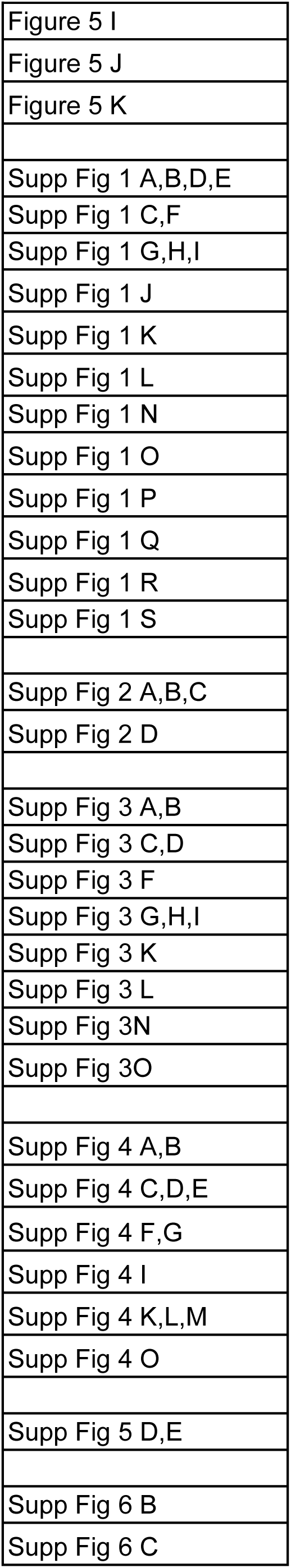
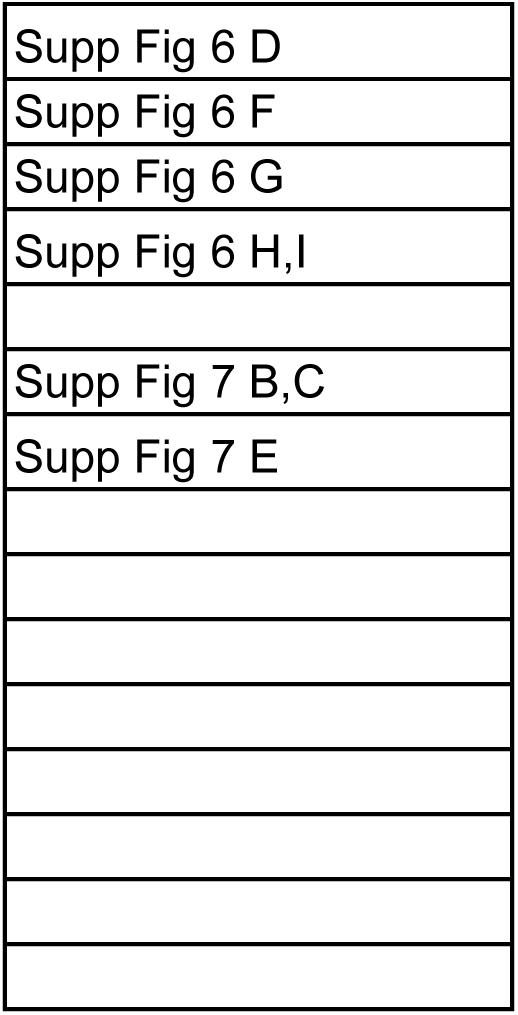

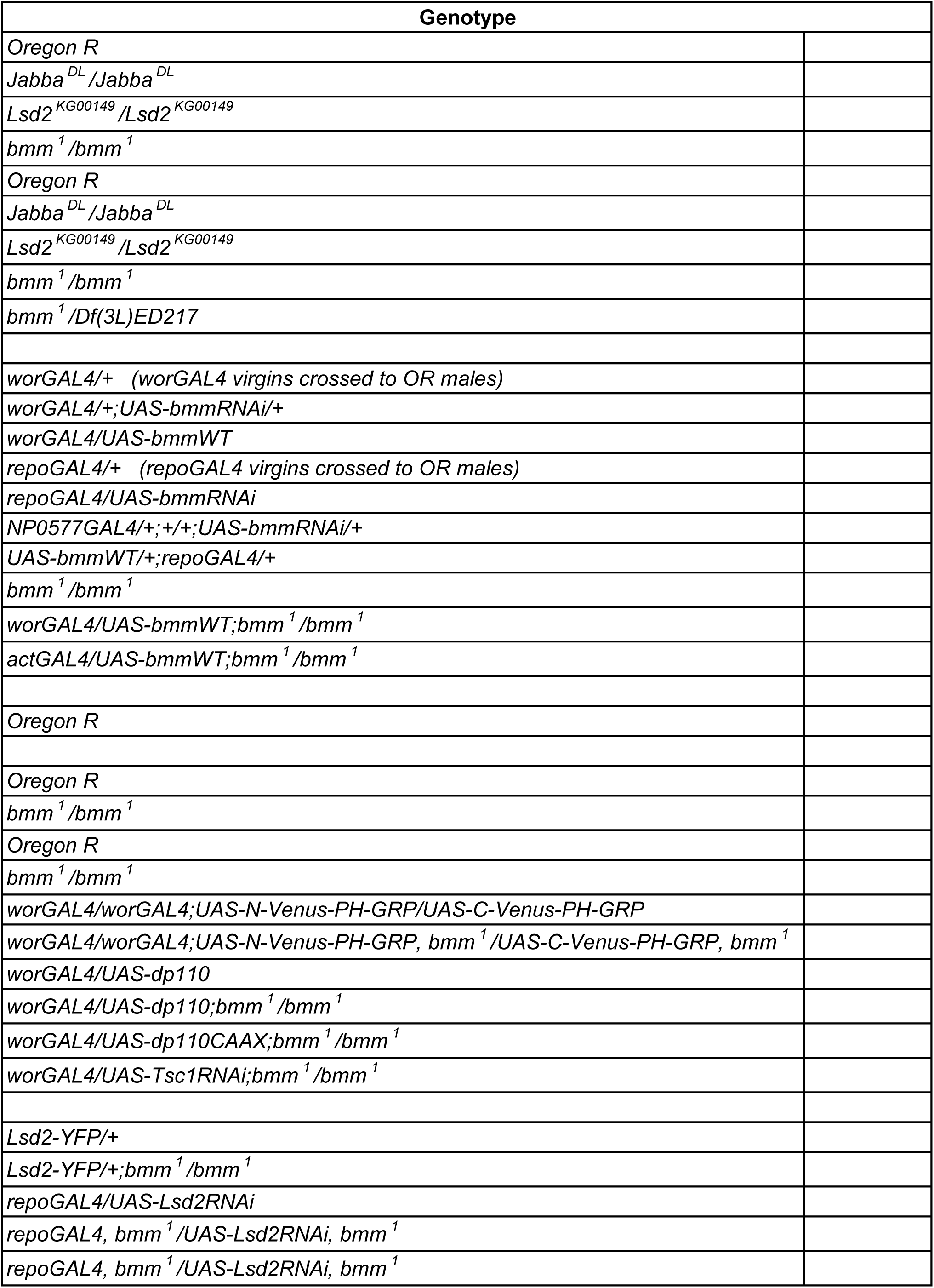
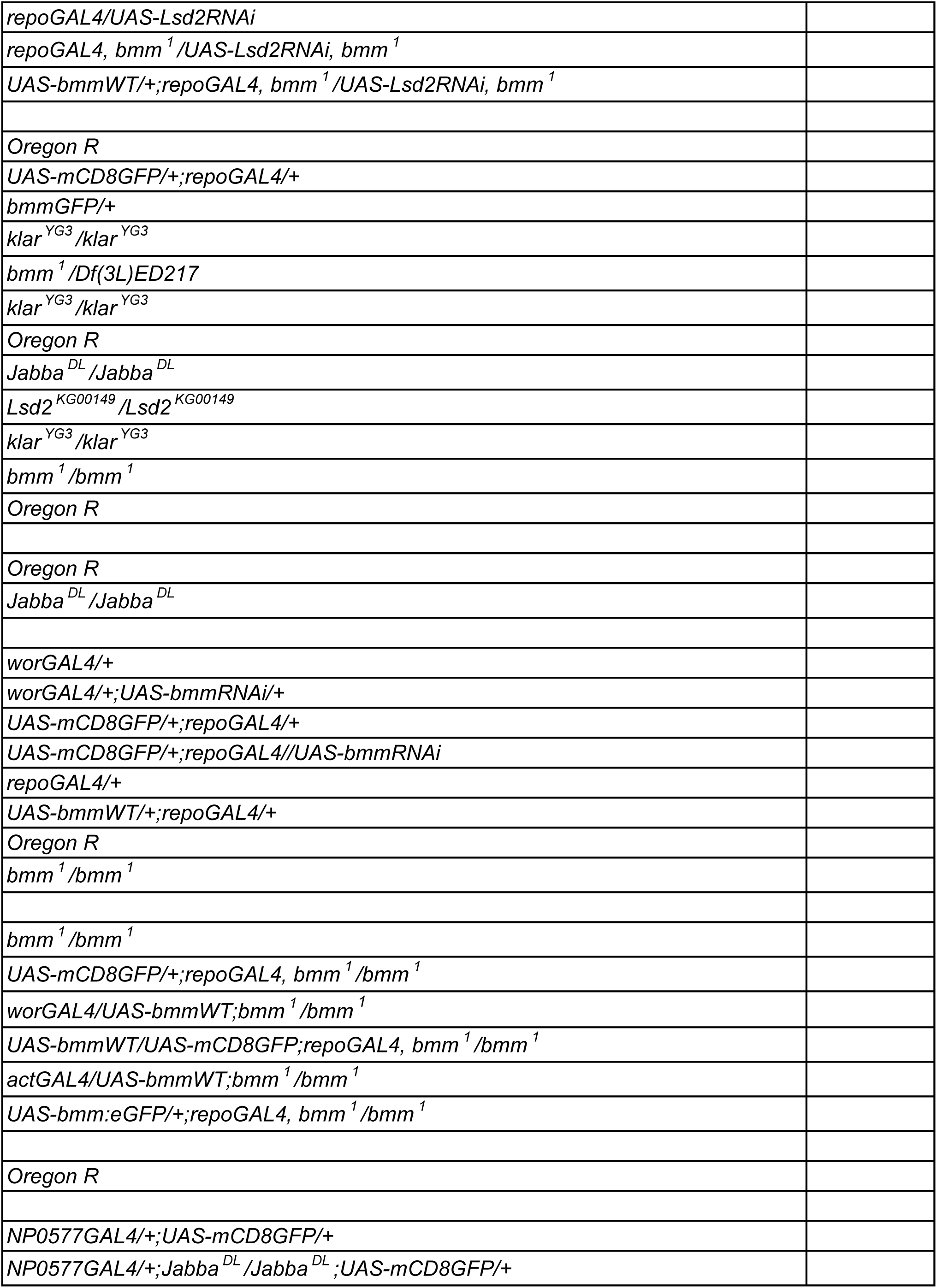
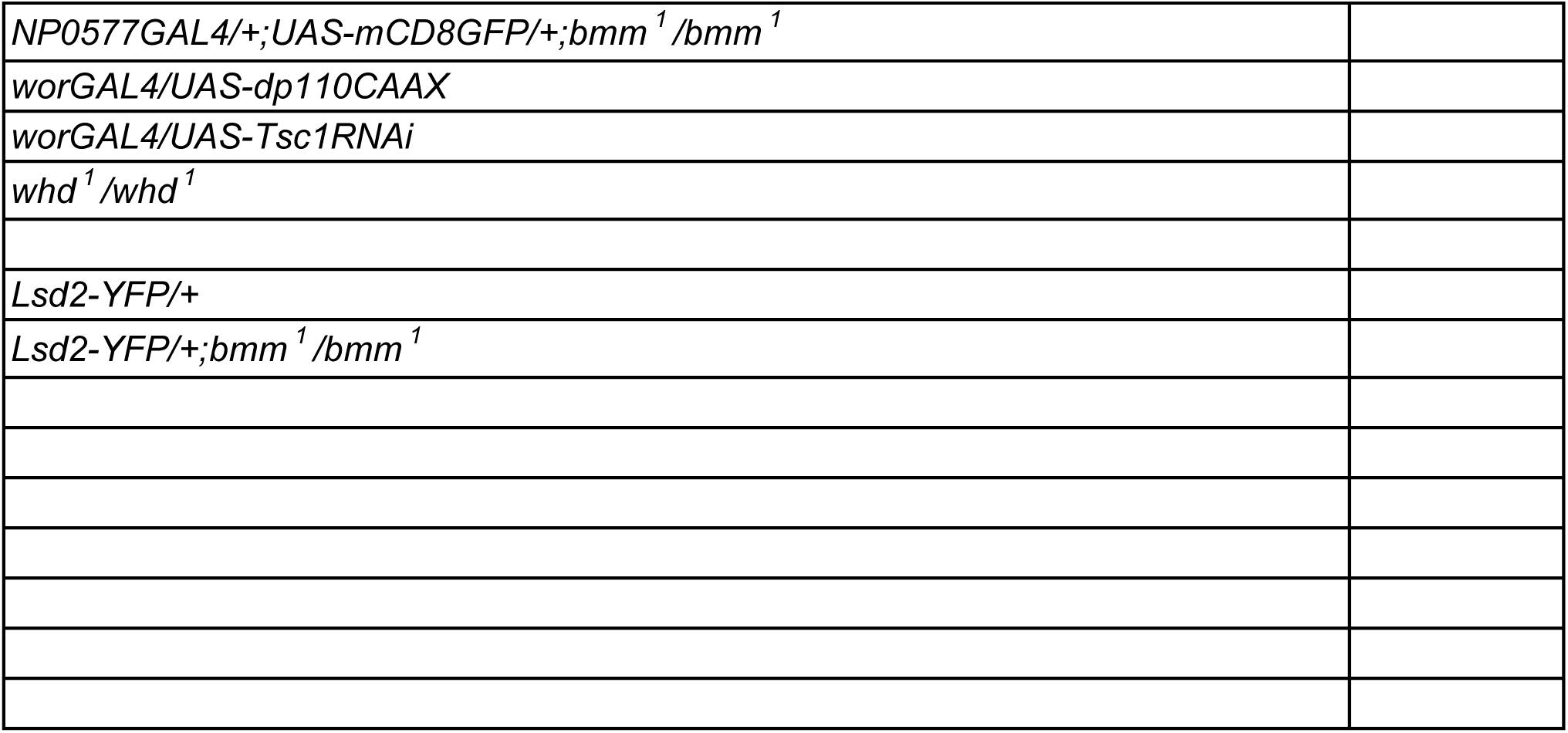

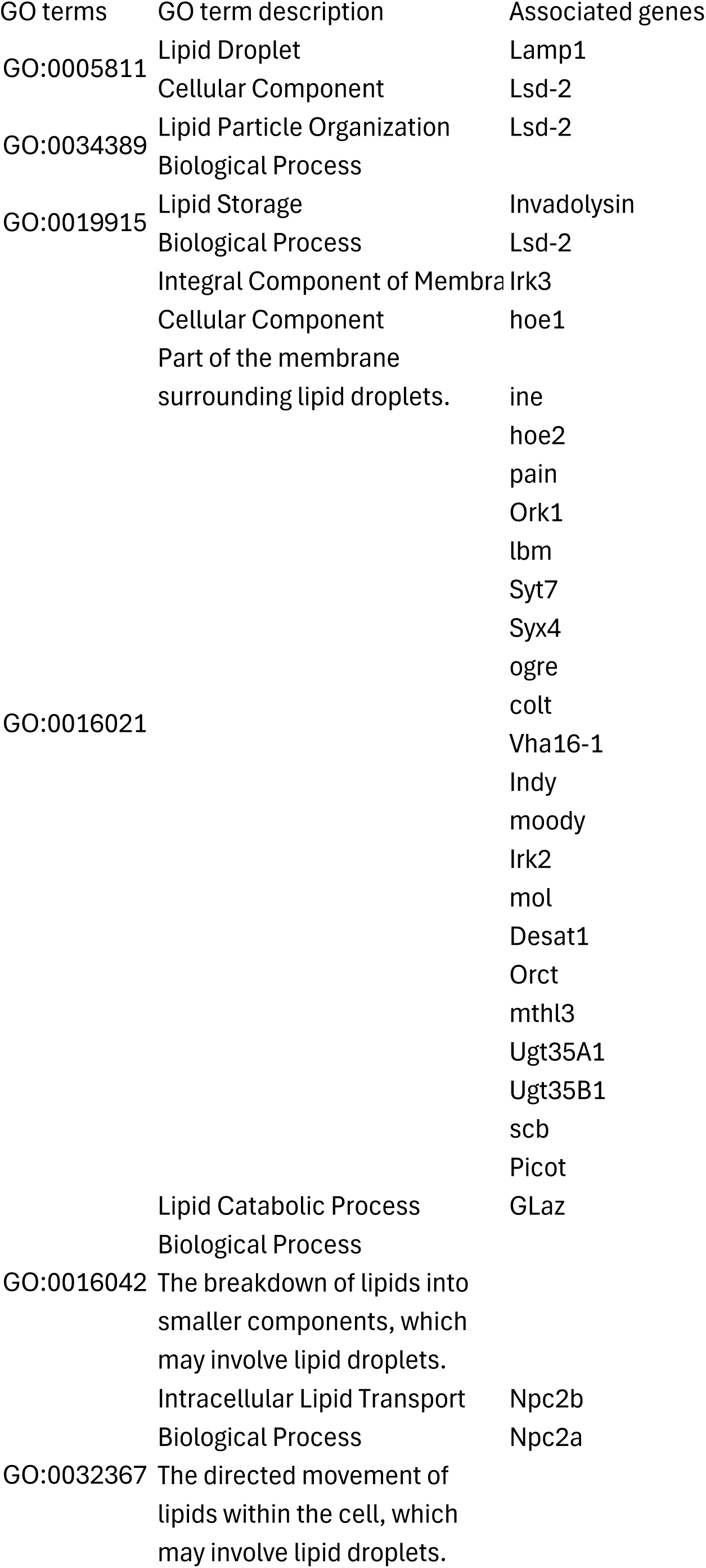
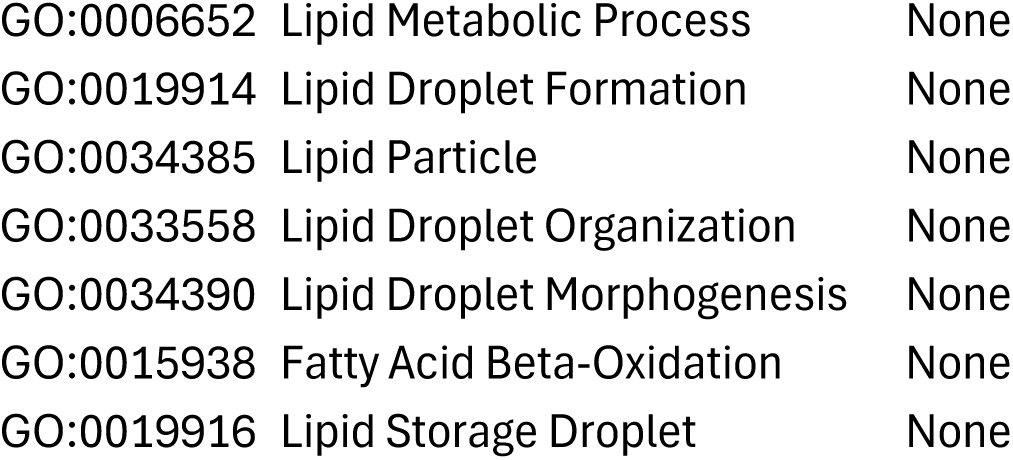

